# FINDER: An automated software package to annotate eukaryotic genes from RNA-Seq data and associated protein sequences

**DOI:** 10.1101/2021.02.04.429837

**Authors:** Sagnik Banerjee, Priyanka Bhandary, Margaret Woodhouse, Taner Z. Sen, Roger P. Wise, Carson M. Andorf

## Abstract

**Background:** Gene annotation in eukaryotes is a non-trivial task that requires meticulous analysis of accumulated transcript data. Challenges include transcriptionally active regions of the genome that contain overlapping genes, genes that produce numerous transcripts, transposable elements and numerous diverse sequence repeats. Currently available gene annotation software applications depend on pre-constructed full-length gene sequence assemblies which are not guaranteed to be error-free. The origins of these sequences are often uncertain, making it difficult to identify and rectify errors in them. This hinders the creation of an accurate and holistic representation of the transcriptomic landscape across multiple tissue types and experimental conditions. Therefore, to gauge the extent of diversity in gene structures, a comprehensive analysis of genome-wide expression data is imperative.

**Results:** We present FINDER, a fully automated computational tool that optimizes the entire process of annotating genes and transcript structures. Unlike current state-of-the-art pipelines, FINDER automates the RNA-Seq pre-processing step by working directly with raw sequence reads and optimizes gene prediction from BRAKER2 by supplementing these reads with associated proteins. The FINDER pipeline (1) reports transcripts and recognizes genes that are expressed under specific conditions, (2) generates all possible alternatively spliced transcripts from expressed RNA-Seq data, (3) analyzes read coverage patterns to modify existing transcript models and create new ones, and (4) scores genes as high- or low-confidence based on the available evidence across multiple datasets. We demonstrate the ability of FINDER to automatically annotate a diverse pool of genomes from eight species.

**Conclusions:** FINDER takes a completely automated approach to annotate genes directly from raw expression data. It is capable of processing eukaryotic genomes of all sizes and requires no manual supervision – ideal for bench researchers with limited experience in handling computational tools.

## Background

Recent advances in sequencing technology enable the construction of chromosomal-level assemblies for even non-model organisms. As of December 2020, genomes of 16,108 eukaryotes, 295,784 prokaryotes, 41,936 viruses, 26,079 plasmids and 17,820 organelles are sequenced and available through GenBank [1], a considerable increase over the 1,500 sequences reported two decades ago (see Additional File 1: Figure S1). Therefore, to annotate the ever-rising number of genome sequences, annotation software applications need to be fast, accurate, and designed to handle large amounts of expression data to facilitate discovery of novel genes across different conditions [2– 5]. Extensive analysis of this available data is the key to achieving exhaustive gene discovery by analyzing samples from multiple tissues and conditions, obviating the need for additional sequencing.

Genome annotation is the process of identifying transcriptionally active regions of the genome and defining gene structures. Decoding the correct structures of genes is essential since several downstream applications rely on accurate annotations: detecting interactions between proteins [6–14], identifying post-translational modifications [15– 23], mining effectors [24–28], and determining protein structure [29–32]. Although we have seen a significant improvement in genome sequencing technology, annotation methods continue to underperform [33, 34]. Obtaining accurate gene annotations is challenging, especially in recently sequenced non-model organisms. The presence of sequences exchanged through horizontal gene transfer in such genomes and the existence of fragmented assemblies make it difficult to predict gene structures [35]. Multiple groups working on the same species have different and oftentimes conflicting annotations that are difficult to merge into a common consensus.

The early 2000s saw initial genome annotation attempts with the introduction of PASA [36], which was developed to map full-length transcripts and Expressed Sequence Tags (ESTs) in order to annotate genomes. In parallel, FGENESH [37, 38], GeneGenerator [39], mGene [40] and GeneSeqer [41] were introduced which predicted gene structures directly from genome sequence. Tools such as MAKER [42–45] and PASA [36] closely depend on pre-assembled full-length transcripts to generate annotations. ESTs and/or *de novo* assembled transcriptomes have been often provided as inputs to these tools to generate annotations [46–52]. Transcripts constructed via *de novo* [53–57] or genome-guided [58–63] approaches are sensitive to the nature of the assembler and its parameter settings. Such assemblers report sequences that are highly similar to one another, making the process of sifting the correct assemblies from artefacts difficult. This issue is moderately mitigated by BRAKER2 [64, 65], which uses read splice information instead of full-length assemblies to predict gene structures and has been shown to perform better than *de novo* approaches [66]. BRAKER2 entails a round of unsupervised gene predictions using GeneMark-ET [67] generating *ab-initio* gene predictions followed by a second round of training by AUGUSTUS [68] using a subset of the gene models created by GeneMark-ET [64]. All variations of MAKER (MAKER, MAKER2 and MAKER-P) use a combination of AUGUSTUS [68] and SNAP [69] to generate gene predictions. Unlike BRAKER2 or PASA, users need to run MAKER for multiple rounds to improve annotation. With no standard technique to optimize the number of rounds, users often undertake a trial-and-error approach to decide what data is supplied to MAKER in each execution round. These unguided choices can create different annotations based on the same data sets. Thus, current approaches report either incomplete genes and/or derive annotations that are missing alternatively spliced transcripts.

To overcome the drawbacks described above, we developed FINDER, a new, automated annotation pipeline that downloads RNA-Seq data from NCBI SRA [70], conducts genome-guided assembly of short reads, predicts gene structure, and annotates genes. FINDER annotates both untranslated and coding regions of genes, categorizes transcripts based on the tissue/conditions where they are expressed, and outputs a complete set of alternatively spliced transcripts. FINDER analyzes the spatial expression profile of each transcript to redefine its boundaries and/or even create newer transcripts and employs an optimized strategy to locate transcripts housing micro-exons. Finally, gene models predicted by BRAKER2 are incorporated into the annotation along with assemblies generated by PsiCLASS [63]. We show that FINDER outperforms state-of-the-art annotation tools in constructing accurate gene structures, when executed with the same expression data.

## Implementation

The detailed workflow of FINDER is outlined in **Fig. 1**. The pipeline accepts metadata via a comma-separated values (csv) file (**see Additional file 2: Table S1**). Users can verify the input data using the `verifyInputsToFINDER` utility (Please check section 1.5.1 of Additional file 9). Both single-end and paired-end data are accepted. The pipeline automatically downloads RNA-Seq data from NCBI SRA or the samples can be accessed locally. Multiple rounds of alignment are conducted using STAR [71, 72] with short reads, thus ensuring the capture of tissue-specific splice junctions and ultimately generates the most comprehensive set of alternatively spliced transcripts. FINDER uses PsiCLASS [63] to generate transcripts both at the tissue level and consolidates them to produce a consensus annotation. It employs change-point detection (CPD) using coverage data to polish intron/exon boundaries if needed. Polished transcripts are then supplied to GeneMarkS-T [73] to predict protein coding regions. In addition to constructing genes from expression data, FINDER uses BRAKER2 [65] to predict genes *de novo*. Finally, gene models are assigned scores that reflect the confidence of prediction and evidence across different data sets. Throughout the pipeline run, intermediate temporary data is removed to optimize space usage. Proper logging of executions is implemented through ruffus [74].

**Fig 1.**
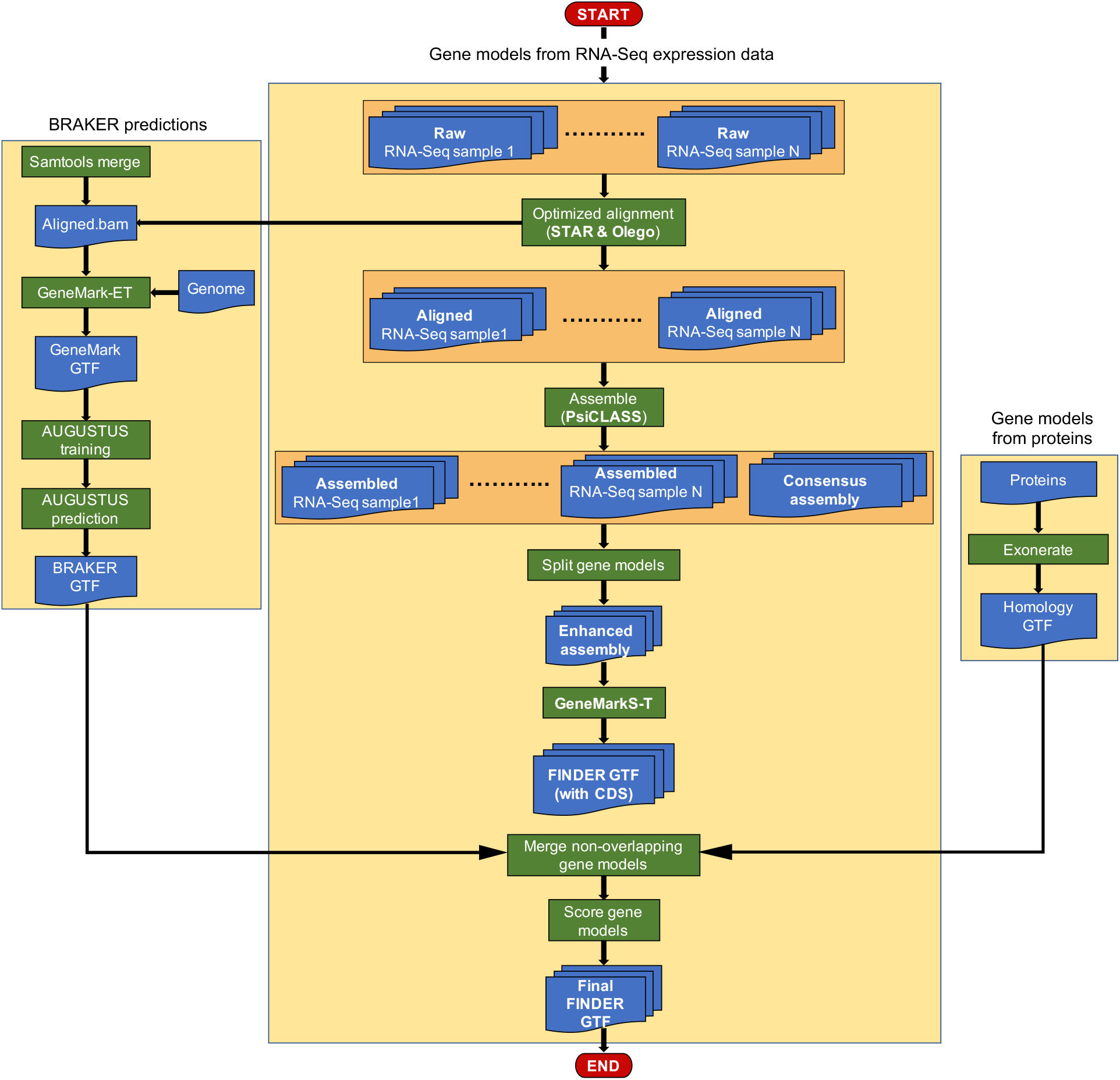
FINDER workflow. FINDER assembles short reads from RNA-Seq expression data, collected from multiple tissues and conditions, to generate full-length transcripts using PsiCLASS. Short read coverage profile is used to polish the structure of the transcripts to enhance the quality of annotation. GeneMarkS-T is used to predict coding regions of the transcripts. Gene models predicted by BRAKER2 and models obtained by mapping proteins are added to the gene models constructed from RNA-Seq data. Additionally, FINDER outputs the tissues where each transcript is expressed allowing users to work with tissue-specific transcripts. FINDER categorizes transcripts into two confidence levels depending on the available supporting evidence and depth of coverage.

### Read alignments to the genome

Reads from each sample are aligned to the genome using STAR [72]. FINDER accepts the location of the genomic STAR indices. If indices are not provided, then FINDER will generate them locally. FINDER implements multiple strategies to detect as many correct splice-junctions as possible. Several studies use a multi-step approach where splice junctions are detected in the first pass and then those junctions are used to guide the alignments in future passes [75, 76]. FINDER employs a similar strategy to align reads and obtain the most confident splice junctions in each tissue type and/or condition by conducting mapping in four passes (Please check section 1.3 of Additional file 9 for more details).

### Annotating transcripts with micro-exons

Certain genes in eukaryotes have micro-exons (i.e., exons with fewer than 50 nucleotides) [77–80] which impart important biological properties both in plants [81–85] and animals [86–90]. FINDER uses OLego [91] to map the reads which were reported unmapped by STAR, because OLego optimizes micro-exon sensitivity by checking intron signatures when no hits of seed sequences (~14 nt) are found. It is configured to align reads to exons of minimum length 2, with a minimum and maximum intron size of 20 and 10K respectively.

### Generating exon-exon transcript structure annotation with PsiCLASS

Alignments reported by STAR and OLego are combined and provided as input to PsiCLASS [63]. Unlike traditional assemblers, PsiCLASS accepts alignments from multiple samples at the same time. It generates annotations for each sample and one consolidated gene annotation for all the samples. FINDER runs PsiCLASS with the -- bamGroup option enabled which instructs PsiCLASS to preserve tissue/condition specific features. It is a fast meta-assembler generating 350 samples of output in less than three hours while running on 30 cores and consumes less than 50 GB of memory.

### Polishing gene structures to optimize gene discovery

Gene structure annotations reported by PsiCLASS were polished to generate the best assemblies. Annotations generated by assemblers often have three kinds of errors that impact accuracy: (1) presence of redundant transcripts that are proper subsets of other transcripts, (2) multiple transcripts on the same strand merged into one, and (3) transcripts with ill-defined exon boundaries. Most assemblers ignore such cases to boost the speed of operation. Developing solutions to deal with these kinds of errors increases the number of correct structural annotations thereby improving downstream analysis.

FINDER uses different algorithmic and statistical approaches to deal with the above cases. To eliminate redundant transcripts, exon-intron structure of all transcripts is compared with each other to retain only unique transcripts. Even though eukaryotes possess large genomes, certain genes/transcripts are closely packed and are overlapping (Fig. 2). Reads originating from one of those genes often map to nearby overlapping genes making the task of distinctly recognizing the transcripts very challenging.

**Fig 2.**
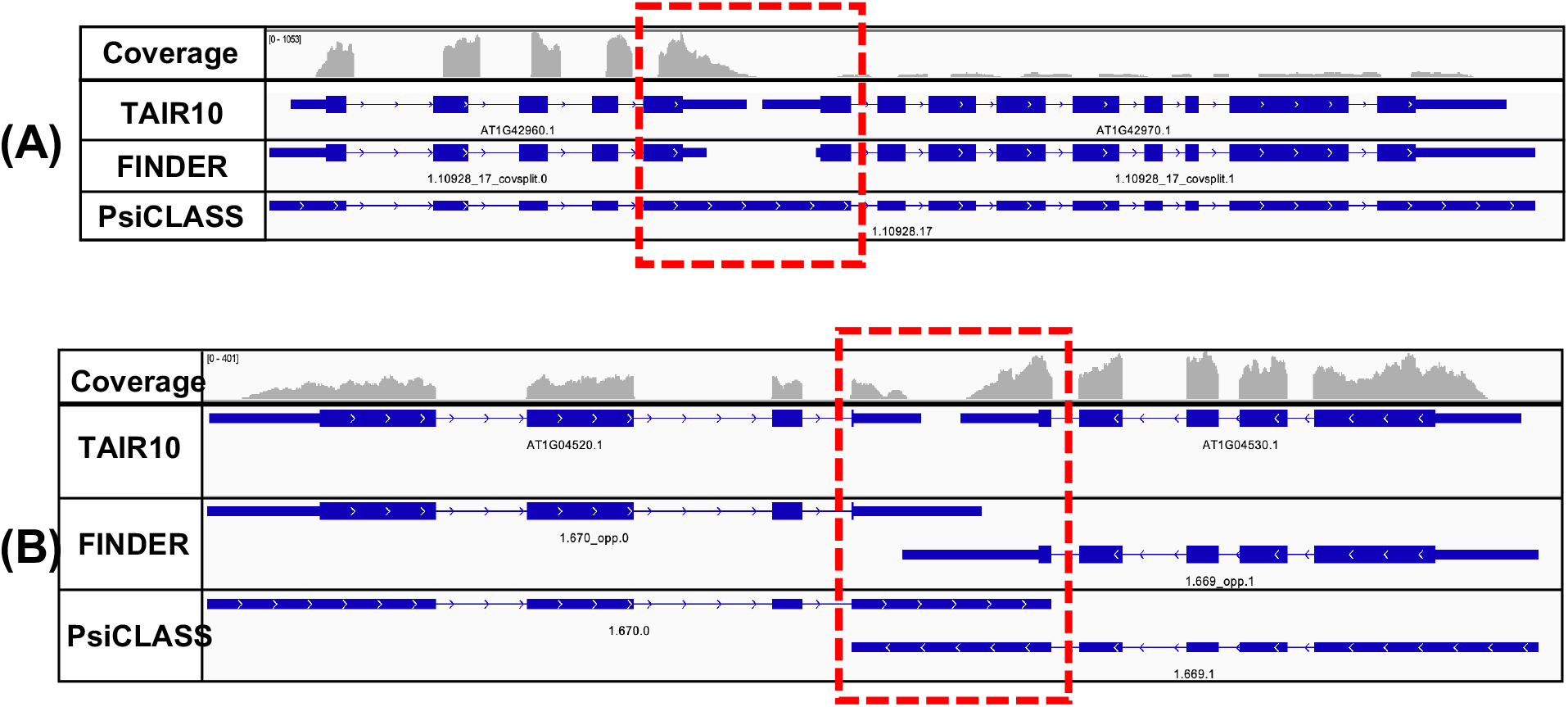
FINDER implements changepoint analysis of read coverages to modify existing gene models and/or generate new ones. Changepoint analysis is a statistical technique to assess alterations in trends over time. The same approach has been used to analyze read coverage patterns of a genome, where the data is distributed spatially. (A) Two *Arabidopsis thaliana* genes AT1G42960.1 and AT1G42970.1 are present within 50 base pairs of each other on the positive strand. Reads originating from the end exons of either genes bleed into each other resulting in PsiCLASS to merge the two gene models. Changepoint analysis recognizes the fall the read coverage and reports a position within the exon where the trough exists. This information is used to split up the gene models. (B) A similar issue exists with closely spaced genes residing on opposite strands. The end exons (highlighted with a red box) for a transcript extend up to the nearest intron of the adjacent transcript. Changepoint analysis is used to determine the actual end/start of transcript based on the read coverage.

FINDER is configured to use changepoint detection (CPD) analysis to detect the descent in read coverage at the junction of two overlapping transcripts. Statistical CPD is a procedure to detect changes in the probability distribution of a stochastic process. Typically, CPD is widely used to detect changes in time series [92–96], but can be extended to other applications as well [97, 98]. We have found that even though CPD was developed under the assumption of normality, it can also be used where normality is violated.

In the first step in FINDER’s CPD, short read alignments to the genome are converted into number of read counts per nucleotide using bedtools [99]. A custom python script is used to transfer the per nucleotide coverage data from the genome to the transcriptome reported by PsiCLASS. Each internal exon is considered as a potential site for the presence of changepoints if there exist premature stop codons in all the three frame translations. CPD only considers exons that have a high chance of housing a changepoint, thereby reducing duration of operation. The coverage pattern of each exon is probed to detect changepoints. The data has been modeled using an exponential distribution, and binary segmentation has been used to determines the changepoints in the exonic coverage using the ‘changepoints’ package [100]. Read coverage of exons mimics a time series where each nucleotide position of an exon can be assumed to be a single unit of time. Coverage patterns of exons, suspected to be merged, contain a characteristic depression in the signal to split the gene models (**Fig. 2A**). Overlapping transcripts on opposite strands sometimes share a common exon (**Fig. 2B**). This negatively impacts precision since the boundaries of the predicted transcript exceed the boundaries of the transcript in the reference annotation. FINDER trims the transcript boundaries, using the changepoints, to better model the RNA-Seq coverage (**Fig. 2B**). These strategies improve the annotation by increasing the transcript F1 scores (**Table 1**).

**Table 1:** Sensitivity, Specificity and F1 scores of transcripts generated by multiple gene annotation pipelines for three model organisms – *Arabidopsis thaliana, Oryza sativa* and *Zea mays*

|  | *Arabidopsis thaliana* |  |  |  |  |  |  | *Oryza sativa* |  |  |  |  |  |  | *Zea mays (NCBI)* |  |  |  |  |  |
| --- | --- | --- | --- | --- | --- | --- | --- | --- | --- | --- | --- | --- | --- | --- | --- | --- | --- | --- | --- | --- |
|  | BRAKER2 | MAKER2 | PASA | FINDER | FINDER+BRAKER2 | FINDER+BRAKER2+PROTEIN |  | BRAKER2 | MAKER2 | PASA | FINDER | FINDER+BRAKER2 | FINDER+BRAKER2+PROTEIN |  | BRAKER2 | MAKER2 | PASA | FINDER | FINDER+BRAKER2 | FINDER+BRAKER2+PROTEIN |
| Base Specificity | 91.08 | 74.87 | 62.71 | 74.46 | 75.01 | 75.04 |  | 57.63 | 52.67 | 36.77 | 42.46 | 42.6 | 42.75 |  | 6.52 | 45.22 | 60.39 | 62.97 | 62.96 | 62.92 |
| Base Sensitivity | 60.27 | 52.55 | 69.41 | 71.45 | 74.14 | 74.23 |  | 36.85 | 40.17 | 59.19 | 61.43 | 61.83 | 62.37 |  | 50.43 | 54.38 | 66.7 | 72.1 | 72.15 | 72.16 |
| Base F1 score | 72.54 | 61.76 | 65.89 | 72.92 | 74.57 | 74.63 |  | 44.95 | 45.58 | 45.36 | 50.21 | 50.44 | 50.73 |  | 11.55 | 49.38 | 63.39 | 67.23 | 67.24 | 67.22 |
| Exon Specificity | 80.28 | 95.35 | 90.74 | 91.79 | 91.43 | 91.43 |  | 38.4 | 72.31 | 67.37 | 67.67 | 67.5 | 67.62 |  | 15.96 | 65.14 | 84.26 | 79.71 | 79.59 | 79.57 |
| Exon Sensitivity | 73.29 | 55.04 | 67.49 | 69.93 | 71.62 | 71.7 |  | 53.74 | 50.59 | 63.99 | 64.81 | 65.22 | 66.06 |  | 65.65 | 61.84 | 71.84 | 74.38 | 74.4 | 74.4 |
| Exon F1 score | 76.63 | 69.79 | 77.41 | 79.38 | 80.32 | 80.37 |  | 44.79 | 59.53 | 65.64 | 66.21 | 66.34 | 66.83 |  | 25.68 | 63.45 | 77.56 | 76.95 | 76.91 | 76.9 |
| Intron Specificity | 86 | 98.26 | 95.98 | 96.55 | 96.26 | 96.26 |  | 52.8 | 76.87 | 72.36 | 73.24 | 73.19 | 73.26 |  | 21.77 | 74.41 | 88.45 | 84.86 | 84.78 | 84.76 |
| Intron Sensitivity | 84.25 | 60.64 | 76.11 | 76.51 | 78.33 | 78.42 |  | 76.78 | 55.77 | 70.44 | 71.03 | 71.52 | 72.34 |  | 80.88 | 64.99 | 75.19 | 78.12 | 78.15 | 78.15 |
| Intron F1 score | 85.12 | 75 | 84.9 | 85.37 | 86.37 | 86.43 |  | 62.57 | 64.64 | 71.39 | 72.12 | 72.35 | 72.8 |  | 34.31 | 69.38 | 81.28 | 81.35 | 81.33 | 81.32 |
| Transcript Specificity | 49.91 | 76.9 | 55.21 | 60.04 | 59.82 | 59.82 |  | 12.71 | 40.13 | 23.17 | 24.54 | 24.59 | 24.54 |  | 2.75 | 32.67 | 48.6 | 44.78 | 44.84 | 44.78 |
| Transcript Sensitivity | 30.26 | 21.74 | 28.62 | 37.21 | 39.21 | 39.28 |  | 16.35 | 18.01 | 29.51 | 33 | 33.32 | 33.82 |  | 19.57 | 25.01 | 37.94 | 42.25 | 42.3 | 42.31 |
| Transcript F1 score | 37.68 | 33.9 | 37.7 | 45.95 | 47.37 | 47.42 |  | 14.3 | 24.86 | 25.96 | 28.15 | 28.3 | 28.44 |  | 4.82 | 28.33 | 42.61 | 43.48 | 43.53 | 43.51 |
| Gene Specificity | 51.58 | 76.9 | 61.79 | 67.33 | 66.56 | 66.55 |  | 13.46 | 40.13 | 36.81 | 32.79 | 32.76 | 32.38 |  | 2.78 | 32.67 | 55.69 | 52.3 | 52.29 | 52.19 |
| Gene Sensitivity | 50.43 | 37.2 | 44.39 | 57.96 | 61.3 | 61.4 |  | 18.07 | 19.4 | 31.37 | 35.05 | 35.4 | 35.96 |  | 29.02 | 38.14 | 51.57 | 57.35 | 57.42 | 57.43 |
| Gene F1 score | 51 | 50.14 | 51.66 | 62.29 | 63.82 | 63.87 |  | 15.43 | 26.16 | 33.87 | 33.88 | 34.03 | 34.08 |  | 5.07 | 35.19 | 53.55 | 54.71 | 54.74 | 54.68 |

### *De novo* gene prediction from expression data and proteins from closely related species

Certain genes are expressed only under specific tissues and conditions [101]. However, constructing an exhaustive set of genes expressed across all possible tissues and conditions is a daunting task due to the mammoth volume of potential expression data. Hence, approaches that can predict structures of unknown genes using information obtained from known genes are needed. Within the FINDER framework, we used BRAKER2 [65] to predict the structure of protein coding genes. The pipeline is provided with alignment files generated by STAR and an optional, user-provided protein data file. If the previous execution fails, a second execution of BRAKER2 is launched without protein information. Genes predicted by BRAKER2 are compared to the genes obtained from expression data. To prevent too many false positives, predictions made by BRAKER2 are considered high confidence, only if those are supported by expression level or protein level evidence.

### Prediction of coding regions

We leveraged GeneMarkS-T [73] to predict protein-coding regions of genes constructed from expression data. GTF files are first converted to FASTA files using the provided genome. Those FASTA files are supplied to GeneMarkS-T as inputs. GeneMarkS-T outputs coding sequence for the transcripts. CDS annotations are incorporated into the final GTF file by converting the transcriptomic coordinates to genomic coordinates.

### Using proteins to annotate more genes

In addition to RNA-Seq data, FINDER also uses protein data (when provided), in two ways (1) to assess the veracity of the transcript models generated by BRAKER2, and (2) to align those proteins not recognized by BRAKER2 or PsiCLASS. Protein coding genes obtained from expression data and predicted by BRAKER2 are BLASTed [102] to the protein set provided by the user. Proteins not encountering any hits are aligned to the genome using exonerate [103] with a minimum threshold of 90% similarity. These alignments are augmented to the final set of gene predictions. Since these transcripts are obtained solely from proteins, they lack UTR sequences.

### Tissue/condition specific transcripts/gene models

Most eukaryotic genes have multiple isoforms which are derived from alternative transcripts. Expression of different transcripts can occur under different conditions in different tissues at different time points. FINDER compares assembled transcripts from each condition and prints out an association between each transcript and the provided tissue/condition (Additional file 9 section 1.5).

### Scoring gene models

FINDER groups genes into multiple categories based on supporting evidence. Genes that are expressed in RNA-Seq datasets, predicted by BRAKER2, and have protein evidence, are put into the high-confidence gene set. BRAKER2-predicted genes with no evidence of expression and/or proteins are treated as low confidence genes. FINDER expects a soft masked genome since it is a BRAKER2 requirement. Genes which are located in the repeat regions are marked as such and moved to the set of low-confidence genes.

## Results & Discussion

### Choice of species for comparison

We tested the performance of FINDER primarily on three well-annotated plant organisms -*Arabidopsis thaliana* [104], *Oryza sativa* [105–107] and *Zea mays* [108, 109]. The genomes assemblies of these model organisms have been frequently updated and are almost complete with telomere-to-telomere sequences with fewer gaps and unknown nucleotides. In addition, their gene annotations have undergone regular improvement by mining the large number of RNA-Seq datasets available in the literature. Also, The Arabidopsis Information Resource (TAIR) provides a five-star rating system based on available evidence for each gene. Such a system offers a platform to test the quality of gene annotation software. For further evaluation, and to ensure that FINDER is able to annotate a wider range of genome types, we selected the following additional species to test: C*aenorhabditis elegans* [110], *Drosophila melanogaster* [111, 112], *Homo sapiens* [113, 114], *Hordeum vulgare* [115], and *Triticum aestivum* [115– 118]). The genomes of these species range from small (*C. elegans, D. melanogaster, A. thaliana*), medium (*O. sativa*), to large (*H. sapiens, Z. mays, H. vulgare*, and *T. aestivum*). Finally, we evaluated FINDER on three different versions of *Z. mays* annotations – RefSeq [119], AGPv3 [109, 120] and AGPv4 [108, 121].

### Metrics to assess quality of annotation

We used four metrics to compare the quality of annotations generated by each pipeline: 1) Annotation Edit Distance (AED) [42, 43, 122], 2) sensitivity, 3) specificity, and 4) F1 score. Although these metrics could be computed both at the nucleotide- and exon-level we chose to make comparisons at the transcript level since it encompasses bases, exons, and introns. An AED score of 0 indicates complete agreement of the predicted annotation with the reference, and a score of 1 denotes that the reference has not been identified in the annotation. A transcript is considered to be “recognized” only when all its intron definitions agree with at least one transcript from the predicted set. We used the Mikado “compare” utility to compare the predictions with the reference annotations [123]. A highly sensitive annotation is one that can correctly recognize more reference transcripts. A set of annotations has high specificity when it reports minimal incorrect transcripts. For an annotation to be of good quality, both sensitivity and specificity should be high. A balanced metric is the F1 score which is the harmonic mean of sensitivity and specificity. While AED provides a good numeric assessment of how well the ground truth evidence is represented in an annotation, when individually used, it fails to capture the extent to which false positives are reported. Hence, F1 score complements AED since it incorporates both specificity and sensitivity. For evaluation purposes, we assume that the annotations achieved through community efforts are the ground truth and contain no errors.

### FINDER generates more accurate gene models than BRAKER2, MAKER2 and PASA

FINDER leverages expression data to construct transcript models and employs statistical changepoint detection to enhance their structures (see Implementation). Both MAKER2 and PASA were run with transcript sequences reported by PsiCLASS.

To assess FINDER’s performance, we compared the AED scores of transcript models generated by FINDER with those generated by other commonly used annotation methods. As shown in **Fig. 3A, 3D & 3G**, the violin plots for FINDER are broader at the base, indicating a greater number of transcripts with lower AED scores as compared to BRAKER2, MAKER, and PASA. We compared the FINDER AED scores with the AED scores reported by other pipelines using Wilcoxon’s signed rank test (More details in Additional file 9 section 2.5). For all organisms (Fig. 3, **Additional file 1: Figure S2–S5 and Additional file 3: Table S2**), the AED scores reported by FINDER were significantly lower (p_value<0.01) than that of any other pipeline. **Fig. 3C, 3F & 3I**, shows a stacked bar plot to represent the fraction of transcripts in each category of AED values. In all the cases, a higher percentage of transcripts reported by FINDER have lower AED scores (**Additional file 1: Figure S2-S5**). This indicates that FINDER is capable of constructing gene structures that better comply with the reference annotations.

**Fig 3.**
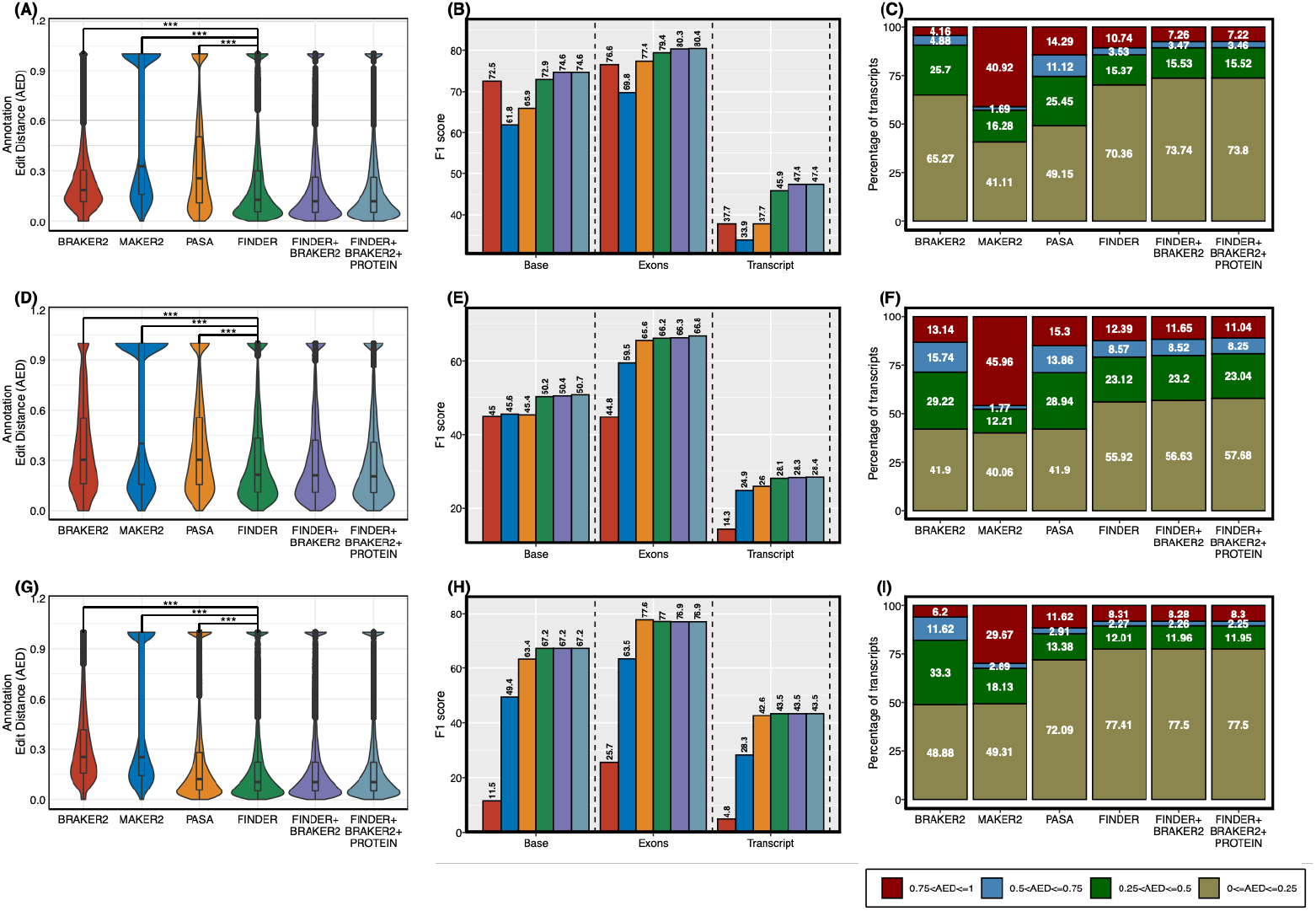
Comparison of performance of predicted annotations in three model species – (A – C) *A. thaliana*, (D – F) *O. sativa* and (G – I) *Z. mays*. Annotation Edit Distance (AED) is an assessment of how well predicted annotations agree with the evidence and was used as a quality control metric. A value of 0 denotes complete agreement of two annotations while a value of 1 denotes that the ‘gold standard’ reference annotation was not detected. Transcripts from ‘gold standard’ reference annotations that are not detected in any of the predicted annotations are removed from analysis. (**A, D & G**) Distribution of AED scores. Violin plots wider at the base indicate high density of annotations with lower AED. FINDER was able to create gene models having lowest AED resulting in a wide base. Gene models generated by FINDER were enhanced by adding predictions made by BRAKER and including protein evidence. Wilcoxon’s signed rank test was used to compare the AED scores between FINDER and other annotating pipelines. The “***” symbol implies that the AED scores of FINDER gene models were significantly lesser (p_value<0.01) than the AED scores of the gene models reported by other pipelines. (**B, E & H**) Bar plot of F1 score of multiple approaches of annotation. Having a high nucleotide F1 (Base F1) or a high exon F1 score is not sufficient to conclude a good annotation. High value of transcript F1 score is indicative of good gene models with high sensitivity and high specificity. (**C, F & I**) Stacked bar plot showing percentage of transcripts in each of the four groups of AEDs. Higher number of transcripts to low AED denotes better annotation. In each of the three species, FINDER was able to generate a higher percentage of transcripts with low AED compared to other techniques of annotation.

High-quality exhaustive annotations predict the fewest false positives thereby boosting the transcript F1 score. The transcript F1 scores of the gene models that were reported by FINDER for *A. thaliana, O. sativa* and *Z. mays* were higher than the models generated by BRAKER2, MAKER, and PASA (**Fig. 3B, 3E & 3H**). This same trend is observed for other tested organisms where FINDER was successful in detecting nucleotides, exons, introns, transcripts and genes (**Table 1, Additional file 1: Figure S2-S5 and Additional file 3: Table S2**). MAKER2 and BRAKER2 registered a high specificity for most of the organisms because fewer transcripts were reported than FINDER. MAKER2 and BRAKER2 also had lower F1 scores, indicating less sensitivity than FINDER. Additionally, we compared the CDS regions of genes reported by FINDER with those of BRAKER2. For most of the organisms, FINDER generated transcript models with a higher F1 score (**Additional file 4: Table S3**). These results show that the better performance of FINDER is ensured not only due to the presence of UTRs but also due to enhanced CDS structure of gene models.

Finally, including BRAKER2 predictions and protein sequences to FINDER enhanced the gene model predictions. About 15% of the gene models reported by BRAKER2, those having high sequence similarity with the provided protein sequences were included in the final annotations (**Table 2**). As shown in **Table 1 and Additional file 5: Table S4**, including evidence at the protein level led to the identification of more genes.

**Table 2:** Improvement in overall gene recognition by adding gene models predicted by BRAKER2 and aligning protein sequences

| Organism | Number of transcript models borrowed from BRAKER | Percentage of transcript models borrowed from BRAKER | Improvement in average annotation score | Number of transcript models from protein alignments | Percentage of transcript models from protein alignments | Improvement in average annotation score |
| --- | --- | --- | --- | --- | --- | --- |
| *Arabidopsis thaliana* | 1692 | 5 | 1.43 | 185 | 0.01 | 0.05 |
| *Oryza sativa* | 5662 | 10 | 0.15 | 440 | 0.01 | 0.15 |
| *Zea mays* | 1061 | 2 | 0.05 | 452 | 0.01 | -0.02 |
| *Caenorhabditis elegans* | 4807 | 18 | 0.48 | 389 | 0.01 | 0.58 |
| *Drosophila melanogaster* | 2421 | 9 | 0.44 | 481 | 0.02 | 0.22 |
| *Homo sapiens* | 5776 | 16 | 0.05 | 229 | 0.01 | 0.15 |
| *Hordeum vulgare* | 1065 | 3 | 0.01 | 19 | 0 | -0.57 |

Unlike BRAKER2, FINDER does not assume a homogeneous nucleotide composition of the genome [124]. FINDER outperforms BRAKER2 while constructing gene models in complex organisms like *H. sapiens, H. vulgare*, and *Z. mays* since assemblers generating transcriptomes from alignments do not require a genome to possess homogeneous nucleotide composition.

FINDER in itself is restricted to annotate genes only in regions of the genome that are transcriptionally active. Recognizing that BRAKER2, being a gene predictor, can construct gene models in transcriptionally silent regions of the genome, FINDER is designed to incorporate the gene models predicted by BRAKER2 into the final annotations.

### Distinct gene groups are accurately annotated with FINDER

Although eukaryotic genes differ from one another in terms of location, structure and the isoforms they encode, most annotation pipelines annotate and evaluate gene predictions with a global and uniform approach. The problem arises when these variances prompt each pipeline to perform differently on dissimilar groups of genes. To avoid this pitfall, we created groups of genes and transcripts based on various criteria (**Table 3)** and compared the performance of FINDER with BRAKER2, MAKER, and PASA for each of these sets.

**Table 3.** Classification of gene models into different groups based on their relative location to other genes, number of isoforms and other criteria.

|  | **Name** | **Description** |
| --- | --- | --- |
| **Group 1** | Uni-exon transcripts | Transcripts having a single exon and no introns |
| **Group 2** | Transcripts without UTRs | Transcripts missing either the 5' or the 3' UTR sequence |
| **Group 3** | Transcripts with UTRs | Transcripts having both UTRs |
| **Group 4** | Transcripts with micro-exons | Transcripts where at least one exon has length less than 50 nucleotides |
| **Group 5** | Transcripts with long introns | Transcripts where at least one intron has a length greater than 10,000 bp |
| **Group 6** | Closely placed transcripts on same strand | Transcripts on the same strand having less than 250 nucleotides between each other |
| **Group 7** | Closely placed transcripts on opposite strand | Transcripts on the opposite strands having less than 250 nucleotides between each other |
| **Group 8** | Multi transcript gene | Transcripts of a gene that have multiple transcripts |
| **Group 9** | Single transcript gene | Transcripts of a gene that have single transcript |

On the set of UTR-containing transcripts, FINDER reported the best transcript F1 scores (**Fig. 4, Additional file 1: Figure S6, S7**). Unlike BRAKER2, FINDER uses GeneMark S/T to predict CDS from the transcript sequences assembled by PsiCLASS and can hence annotate UTR regions. For most of the organisms, BRAKER2 and MAKER2 gene models register a low transcript F1 score in this category of genes. Next, we tested the performance of the annotation pipelines on transcripts that are closely located in the genome. On this set of transcripts, FINDER reported the best F1 transcript score for *A. thaliana, O. sativa*, and *Z. mays* (**Fig. 4**), and comparable scores for *D. melanogaster* (**Additional file 1: Figure S6**), *H. vulgare* (**Additional file 1: Figure S8**), and *C. elegans* (**Additional file 1: Figure S7**) with BRAKER2. Most eukaryotic genes have multiple isoforms which differ from one another by their exon-intron definition. Splice sites and coverage information provides clues to construct such alternatively spliced transcripts. We selected genes with more than one transcript to check how well each annotation pipeline was able to detect transcript isoforms. For this case, FINDER was able to generate the best transcript structures with the highest transcript F1 score among all the pipelines gene annotation software applications (**Fig. 4 and Additional file 1: Figure S6-S9)**. Surprisingly, BRAKER2 fared poorly in this category despite training with all the detected splice sites from RNA-Seq data. This demonstrates that FINDER is capable of leveraging both intron splice sites and read coverages to report best transcript structures. For *H. sapiens*, PASA was able to generate the best transcript structures across all categories of transcripts. Adding transcripts from BRAKER2 and protein evidence improved the transcript F1 score for all the organisms, signifying the importance of incorporating *de novo* gene models and protein evidence.

**Fig 4.**
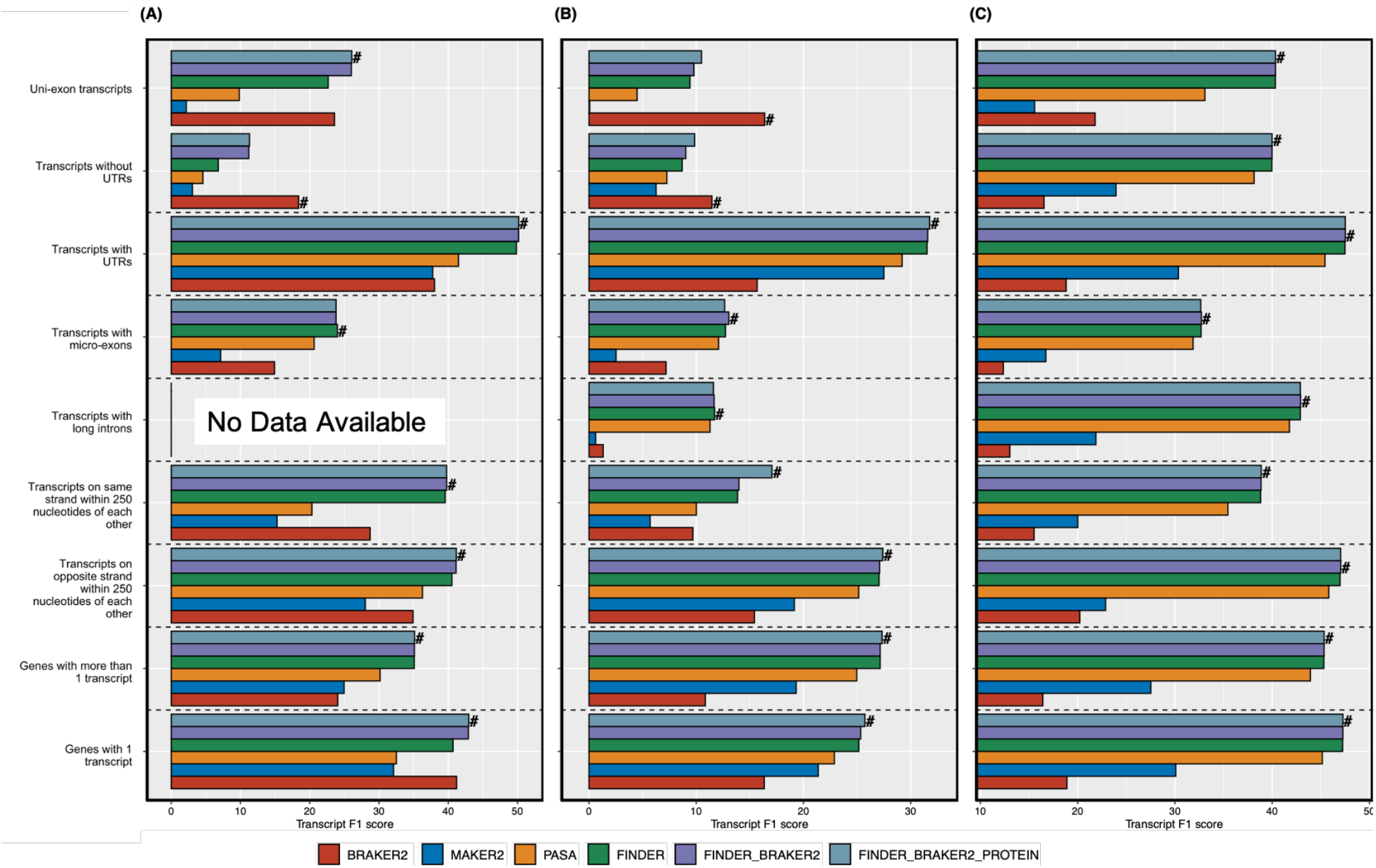
FINDER vs other pipelines on different groups of genes in three model species – (A) *A. thaliana*, (B) *O. sativa* and (C) *Z. mays*. F1 score is the harmonic mean between sensitivity and specificity. Higher F1 score indicates better agreement with the reference transcript models. We created groups of transcripts that have similar characteristics as shown in the y-axis legend. A pool of transcripts was created containing multi-exonic transcript predictions, from each pipeline, that has a complete intron chain match with at least one reference annotation. Mono exonic transcripts were considered if at least 80% of the nucleotides overlap with one reference annotation. Transcript F1 scores, for each of the annotation pipelines, have been plotted as a bar graph. Even though all annotation pipelines are designed to serve the same purpose of annotating genomes, each pipeline adopts a different strategy. Each strategy has its own merits and demerits that lead to better annotation of a certain category of genes. This plot helps understand the performance of each annotation pipeline on different categories. The symbol “#” denotes the best annotator in each gene group.

BRAKER2 generated the best transcript annotation for the set of transcripts with a single exon (**Fig. 4A&B** and **Additional file 1: Figure S6-S9**). Such transcripts, devoid of any introns, are difficult to construct from RNA-Seq alone. Also, the direction of the splice sites infers the direction of a transcript. Without any introns, such a single-exon transcript has to be probed for a CDS sequences’ presence to infer directionality. BRAKER2 was configured to optimally predict only CDS regions of genes, hence, it performs well with the set of transcripts that have missing UTRs for organisms with small and moderate sized genomes (**Fig. 4A&B and Additional file 1: Figure S6-S9)**. The average number of transcripts per gene reported by BRAKER2 is lower than FINDER. While this boosts specificity, it compromises recall since BRAKER2 is not sensitive to detecting alternatively spliced transcripts. Hence, BRAKER2 accomplishes the best F1 score when tested on a set of single-transcript genes but performs poorly on a set of multi-transcript genes (**Fig. 4A&B and Additional file 1: Figure S6-S9**).

### Performance comparison on TAIR’s 5-star System

In order to assess the performance of the annotation pipelines on groups of genes constructed from varying levels of evidence, we used the TAIR10 5-star system. TAIR associates a quality score to each *A. thaliana* transcript based on the evidence used to construct the models, with five stars designating the best evidence and zero stars the least [125]. The three categories with limited evidence (<3 stars) have fewer than 3,000 transcripts each. BRAKER2’s performance, on the genes in these three categories, was slightly better than the rest of the annotation pipelines (**Fig. 5**). The other two categories (five star and four star) have 9,067 and 18,374 transcripts respectively. In both of these categories, FINDER was able to detect more transcripts than any other annotation pipeline. 51.5% and 86.4% of genes in the 5-star and 4-star category respectively were multi-exonic. In both these categories, FINDER correctly constructed more gene models compared to any other annotation pipeline (**Fig. 5**). FINDER reported 80% of the gene models belonging to the 4-star category – 18% more than BRAKER2 (**Fig. 5**). Hence, it is evident from this analysis that FINDER can reconstruct the structures of most of the genes that are well-supported by underlying evidence.

**Fig 5.**
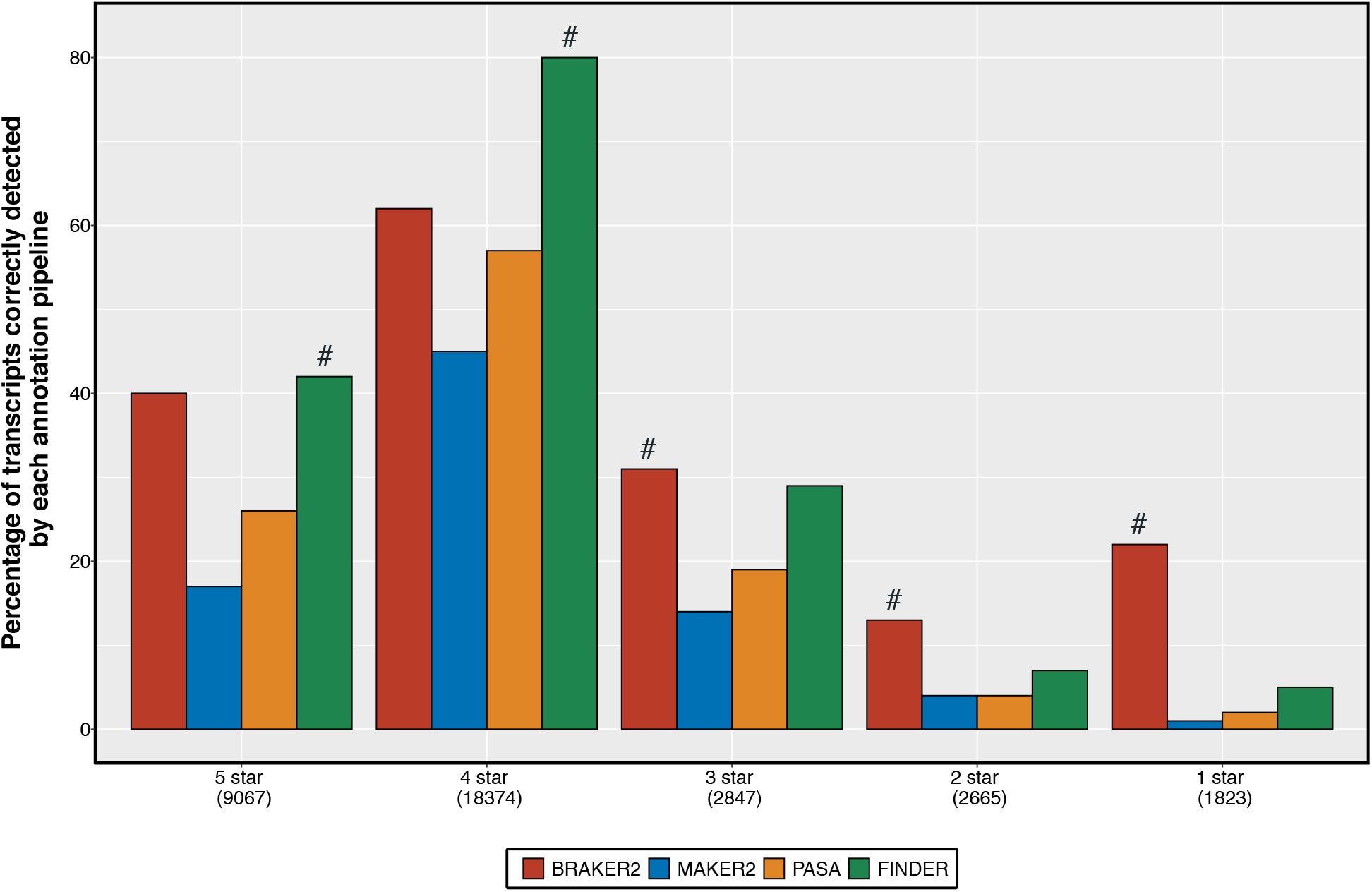
Performance of annotation pipelines on gene groups of *Arabidopsis thaliana* generated by TAIR10. The Arabidopsis Information Resource (TAIR) group has created a quality ranking system to indicate the level of confidence in an annotated gene/transcript. The ranking system has five levels (denoted by stars). Higher number of stars denote the availability of more information to generate the gene structure. Here we display the percentage of transcripts in each category that was identified by a particular annotation pipeline. A high percentage of identified transcripts indicate higher sensitivity and hence a better annotation. The number below each legend in the x-axis denote the number of genes in that respective group. The “#” denotes the predictor which detected the maximum number of transcripts within each group.

### Improving transcript annotations using changepoint analysis

The co-location of multiple overlapping genes on the genome strands makes it difficult to correctly annotate their structures (see Methods **Polishing gene structures to optimize gene discovery**). FINDER employs changepoint detection (CPD) [100] to split the merged transcripts reported by PsiCLASS (**Fig. 2**). To gauge the magnitude of improvement in transcript structures brought about by the application of CPD, we compared the accuracy of the predicted transcriptome before and after implementing CPD based on read coverage. As shown in **Table 4 and Additional file 6: Table S5**, implementing the CPD improved both specificity and sensitivity in organisms with small or medium-sized genomes. In *A. thaliana*, the transcript F1 scores increased from 40.78 to 45.95 (**Table 4 and Additional file 6: Table S5**) and in *C. elegans* it increased from 40 to 50. In large genomes, the improvement was not as significant, mainly because there are only a few genes that overlap with one another.

**Table 4:** Comparison of specificity, sensitivity and F1 scores of transcripts assemblies generated by Strawberry, Scallop, Stringtie, PsiCLASS and FINDER for three model organisms – *A. thaliana, O. sativa* and *Z. mays*.

|  | *Arabidopsis thaliana* |  |  |  |  |  | *Oryza sativa* |  |  |  |  |  | *Zea mays (RefSeq)* |  |  |  |  |
| --- | --- | --- | --- | --- | --- | --- | --- | --- | --- | --- | --- | --- | --- | --- | --- | --- | --- |
|  | STRAWBERRY | SCALLOP | STRINGTIE | PSICLASS | FINDER |  | STRAWBERRY | SCALLOP | STRINGTIE | PSICLASS | FINDER |  | STRAWBERRY | SCALLOP | STRINGTIE | PSICLASS | FINDER |
| Base Specificity | 38.41 | 37.34 | 58.35 | 62.63 | **74.46** |  | 22.6 | 24.23 | 39.83 | 36.64 | **42.46** |  | 30.06 | 29.6 | 49.33 | 55.61 | **62.97** |
| Base Sensitivity | **87.06** | 85.3 | 80.22 | 70.83 | 71.45 |  | **78.2** | 77.64 | 70.87 | 60.27 | 61.43 |  | **81.2** | 79.08 | 76.98 | 70.52 | 72.1 |
| Base F1 score | 53.3 | 51.94 | 67.56 | 66.48 | **72.92** |  | 35.07 | 36.93 | 51 | 45.57 | **50.21** |  | 43.88 | 43.08 | 60.13 | 62.18 | **67.23** |
| Exon Specificity | 43.86 | 70.64 | 74.82 | 89.82 | **91.79** |  | 23.51 | 42.97 | 51.7 | 66.29 | **67.67** |  | 37.18 | 52.37 | 60.33 | 77.76 | **79.71** |
| Exon Sensitivity | **85.3** | 79.67 | 79.29 | 69.54 | 69.93 |  | 79.08 | 76.65 | 75.47 | **65.75** | 64.81 |  | **85.03** | 81.68 | 81.77 | 75.88 | 74.38 |
| Exon F1 score | 57.93 | 74.88 | 76.99 | 78.39 | **79.38** |  | 36.24 | 55.07 | 61.36 | 66.02 | **66.21** |  | 51.74 | 63.82 | 69.43 | 76.81 | **76.95** |
| Intron Specificity | 55.32 | 78.7 | 80.58 | 95.29 | **96.55** |  | 29.13 | 48.56 | 56.5 | 71.41 | **73.24** |  | 43.79 | 56.74 | 64.79 | 82.8 | **84.86** |
| Intron Sensitivity | **92.06** | 89.99 | 87.75 | 77.63 | 76.51 |  | 85.84 | **85.41** | 83.28 | 71.72 | 71.03 |  | **90.19** | 86.69 | 86.05 | 78.99 | 78.12 |
| Intron F1 score | 69.11 | 83.97 | 84.01 | **85.56** | 85.37 |  | 43.5 | 61.92 | 67.32 | 71.56 | **72.12** |  | 58.96 | 68.59 | 73.92 | 80.85 | **81.35** |
| Transcript Specificity | 6.88 | 24.84 | 35.02 | 56.82 | **60.04** |  | 1.59 | 9.03 | 14.26 | 24.22 | **24.54** |  | 6.96 | 17.8 | 26.23 | 43.96 | **44.78** |
| Transcript Sensitivity | 31.68 | 32.19 | 35.23 | 31.8 | **37.21** |  | 26.69 | 29.43 | 31.2 | 32.37 | **33** |  | **48.71** | 46.71 | 47.76 | 42.48 | 42.25 |
| Transcript F1 score | 11.3 | 28.04 | 35.12 | 40.78 | **45.95** |  | 3 | 13.82 | 19.57 | 27.71 | **28.15** |  | 12.18 | 25.78 | 33.86 | 43.21 | **43.48** |
| Gene Specificity | 33.16 | 36.51 | 63.7 | 65.67 | **67.33** |  | 17.61 | 20.16 | 35.69 | **40.39** | 32.79 |  | 21.83 | 26.33 | 47.43 | **54.04** | 52.3 |
| Gene Sensitivity | 44.77 | 46.18 | 50.59 | 49.33 | **57.96** |  | 28.91 | 31.79 | 33.65 | 34.54 | **35.05** |  | 55.2 | 55.85 | 58.03 | 57.22 | **57.35** |
| Gene F1 score | 38.1 | 40.78 | 56.39 | 56.34 | **62.29** |  | 21.89 | 24.67 | 34.64 | **37.24** | 33.88 |  | 31.29 | 35.79 | 52.2 | **55.58** | 54.71 |

### PsiCLASS meta-assembly works better than other approaches

We explored three popularly used software applications for merging transcriptome assemblies – StringTie-merge [76, 126–132], TACO [133–138] and Cuffmerge [139– 144] to combine 116 *A. thaliana* assemblies constructed by StringTie [59], Scallop [61] and Strawberry [60] (Please check section 3 of Additional file 9 for more details). The best assembly was reported by StringTie-merge and was hence used for all other organisms. We compared the accuracy of the consensus transcript models generated by StringTie-merge with the transcript models reported by PsiCLASS [63]. As depicted in **Table 4 and Additional file 6: Table S5**, PsiCLASS generated the best transcript models for all organisms registering the highest transcript F1 score improving upon the StringTie models by up to 15%. Hence, FINDER uses only PsiCLASS to generate assemblies from short-read data.

### Impact of missing untranslated region on annotation of transcripts

Gene transcription is triggered by adherence of a transcription factor in the promoter region of a gene. Promoters are typically located within 1,000 bp upstream of a gene’s transcription start site (TSS) [145–147]. Determining the TSS from sequencing data is best facilitated by RAMPAGE [148, 149] or CAGE-Seq [150], but this data is usually unavailable due to constraints imposed by cost and time. Nevertheless, a good estimate can be obtained from RNA-Seq data by assuming the start coordinates of the assembled genes as the TSS. Thus, researchers often localize their investigation to a section 500-1,000 bp upstream of the assumed TSS [151, 152]. Without 5’ UTR annotation it is impossible to deduce a good approximation of the TSS. This leads to conducting promoter mining in a completely incorrect genome location. To assess the quality of 5’ UTR annotation, we plotted the difference of TSS between the reference genes and the genes reported by BRAKER2 and FINDER using a violin plot (**Fig. 6**). Further, we applied Wilcoxon’s rank-sum test and found that the TSS distances reported by FINDER were significantly less than that of BRAKER2 for *A. thaliana* and *Z. mays*. Interestingly, for *O. sativa*, BRAKER2 generated better gene models for more transcripts. Over 25% of reference gene models in *O. sativa* have no UTRs annotated which is higher compared to 15% UTR-less gene models in *A. thaliana* and *Z. mays*. This result illustrates that more FINDER transcripts have a TSS closer to the evidence as compared to the TSS of the transcripts reported by BRAKER2. This is an expected result since BRAKER2 was configured to annotate only CDS regions of transcripts. **Table 5** highlights the number of transcripts that have better agreement with the reference TSS for FINDER and BRAKER2.

**Table 5.** Use of RNA-Seq evidence to improve annotation of untranslated regions to aid in promoter mining and epigenetic studies

|  | Number of<br>FINDER1<br>transcripts<br>having TSS<br>better than<br>BRAKER2 | Number of<br>BRAKER2<br>transcripts<br>having TSS<br>better than<br>FINDER1 |
| --- | --- | --- |
| *Arabidopsis thaliana* | **15063 (65%)** | 8022 (35%) |
| *Oryza sativa* | **11089 (66%)** | 5762 (34%) |
| *Zea mays* (NCBI) | **20721 (76%)** | 6628 (24%) |
| *Zea mays* (AGPv3) | 7618 (28%) | **19731 (72%)** |
| *Zea mays* (AGPv4) | **18114 (69%)** | 8297 (31%) |
| *Caenorhabditis elegans* | 8681 (33%) | **17730 (67%)** |
| *Drosophila melanogaster* | **10238 (63%)** | 5917 (37%) |
| *Homo sapiens* | **10158 (74%)** | 3486 (26%) |
| *Hordeum vulgare* | **10373 (65%)** | 5607 (35%) |

**Fig 6.**
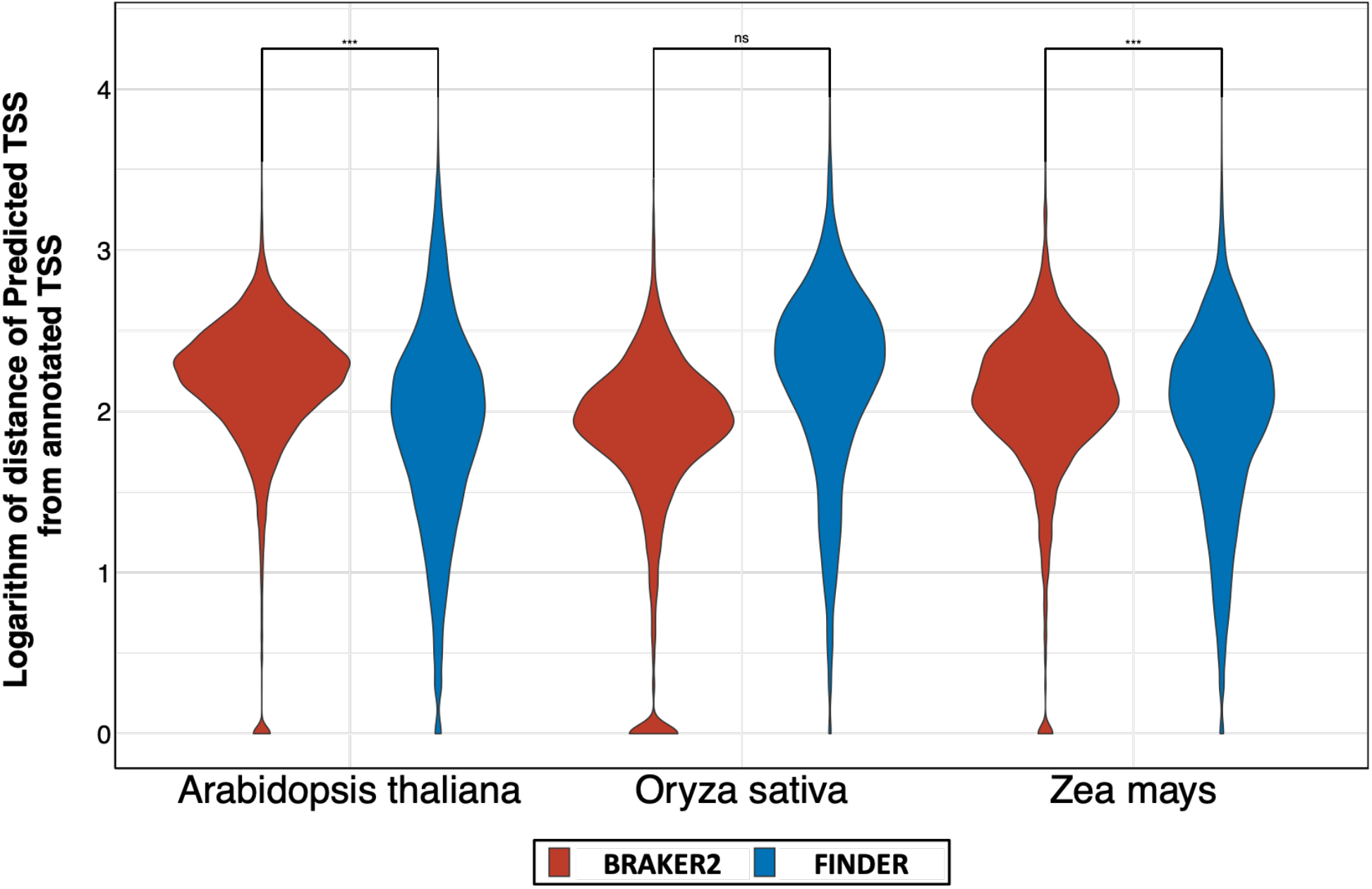
Comparison of distance between transcription start sites of gene models predicted by BRAKER2 and FINDER. Violin plots of the distribution of the distance between the actual transcription start site (TSS) and the predicted transcription start site. In a set of well annotation complete gene structures, a higher fraction of genes is expected to have low deviation from the actual TSS. We considered genes that were reported in either BRAKER or FINDER for this analysis. Wilcoxon’s rank sum test was used to compare the TSS distances between FINDER and BRAKER2. The “***” symbol implies that TSS distance for FINDER gene models was significantly less than BRAKER2 gene models.

### Enhancing ground truth annotations by extending untranslated regions

Official annotations of several model organisms, used as ground truth for this study, contain transcripts with missing UTR sequences. Even though UTRs do not code for proteins, they are relevant segments of a transcript involved in several important biological processes like mRNA translation [153–155], regulation of expression [156– 160]] and a number of diseases [161–165]. In the *A. thaliana* TAIR10 annotations, there are 7,888 transcripts missing either UTR; 50% of these had a rating below 2 stars.

PacBio (Menlo Park, CA) offers long-read sequencing that contain both CDS and UTRs. Therefore, we used the PacBio annotations instead of the incomplete TAIR10 transcripts to assess FINDER’s performance on transcripts that were missing UTRs (Please refer to section 2.6 in Additional file 9 for more details). Out of the 7,888 TAIR10 transcripts with missing UTRs, 113 transcripts were found both in the PacBio data and the 116 short-read RNA-Seq samples. We compared the FINDER annotations against these 113 transcripts. FINDER annotations were able to recall 91.55% of the nucleotides in 113 transcripts of TAIR10 and 97.86% of PacBio transcripts. The specificity of the FINDER annotations is markedly higher with PacBio transcripts (79.67%) compared to TAIR10 transcripts (72.14%). This demonstrates that FINDER enhances and improves upon the existing annotation.

The TRITEX *H. vulgare* annotation (Morex version r2) [115], released by the International Barley Sequencing Consortium (IBSC), is devoid of UTRs. We used FINDER to update and enrich the existing annotations by flanking the CDS region with UTRs on both sides. To verify the accuracy of the gene models reported by FINDER, we used PacBio full-length mRNA sequences derived from a time course of powdery mildew infected barley leaf tissue [166, 167]. A total of 7,352 gene models from IBSC, FINDER, and PacBio had a complete intron-chain match with each other. The gene structures for more than 93% (6,886 out of 7,352) of the FINDER models were improved when compared to PacBio full-length sequences (**Additional file 7: Table cS6**). The highest F1 score achieved was 87.16. This shows that FINDER is capable of constructing accurate gene structures constituting both CDS and UTRs.

### Evaluating performance with different annotations of *Zea may*s

*Z. mays* is an important model organism for crops and has been one of the most studied plants for genetics by researchers in several different fields [168–171]. Genes have been annotated in multiple ways using different kinds of data, resulting in substantial differences in gene structures [120]. Here we compare three alternative annotation sets of *Z. mays* – RefSeq, AGPv3, and AGPv4 and the performance of FINDER surpassed all three approaches. The transcript F1 score for FINDER gene models compared against the NCBI gene models were 43.48, whereas the F1 scores for AGPv3 and AGPv4 were 26.69 and 22.51 respectively. We observed the same trend for other annotation pipelines and reported a higher transcript F1 score for NCBI than the AGP annotations (**Table 1 and Additional file 3: Table S2)**. Hence, FINDER generated high-quality gene structures with high transcript F1 scores for different *Z. mays* annotations.

### Evaluating FINDER on different clades reported by Phylostratr

Genes in each organism can be categorized by their evolutionary history [172, 173]. We used Phylostratr [174] to classify genes into evolutionary strata. Here we present our results on the three model organisms – *A. thaliana, O. sativa*, and *Z. mays*. For all three, FINDER was able to accurately detect more genes in highly populated strata (**Fig. 7**). The performance of FINDER and PASA was comparable in strata with few genes. It was surprising to note that BRAKER2 was unable to identify highly conserved genes (those from the “cellular organisms” strata) since those would be easier to predict than organism specific genes. This demonstrates that FINDER is capable of effectively constructing genes from different evolutionary backgrounds.

**Fig 7.**
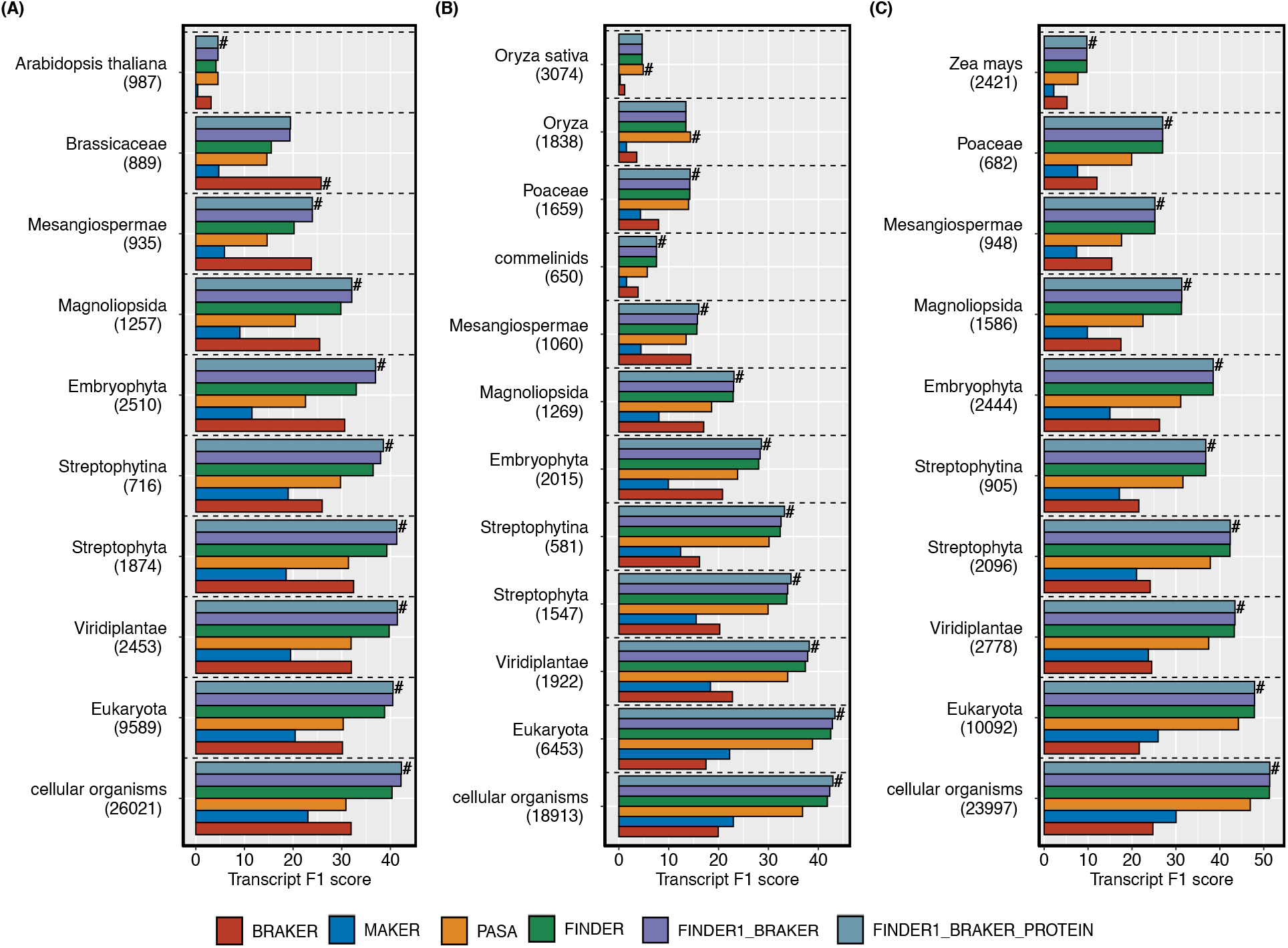
Assessment of annotation pipelines on genes from each phylostrata. – Genes from three model species – (A) *Arabidopsis thaliana*, (B) *Oryza sativa* and (C) *Zea mays*, were allocated into evolutionary classes using Phylostratr. The number of genes correctly constructed by each pipeline was computed and plotted as a bar graph. Numbers below each stratum indicate the number of genes allocated to that strata. Strata having genes fewer than 500 are not shown in the graph.

### FINDER constructs gene models for polyploid genomes

Being a general-purpose genome annotator, in addition to diploid organisms, FINDER can annotate the genomes of polyploid organisms. We generated gene structures of *Triticum aestivum*, a hexaploid with 120,744 annotated genes and 146,597 transcripts [115]. FINDER was able to detect 48,129 transcripts (39.9%). Out of the 130,582 transcripts predicted by FINDER, 48,104 (36.83%) matched perfectly with at least one reference annotation.

## Conclusion

Identifying genes on chromosomes and deducing their structures from a plethora of evidence has been undertaken in multiple ways, with each method having advantages and disadvantages. Herein, we propose FINDER – an entirely automated, general-purpose pipeline to annotate genes in eukaryotic genomes. FINDER (1) implements an optimized mapping strategy that reduces the number of spurious mappings, (2) produces complete full-length transcripts comprising UTRs while identifying transcripts with micro-exons, (3) employs statistical CPD to modify gene boundaries and construct new genes, (4) reports more alternatively spliced transcripts as compared to other state-of-the-art annotation pipelines, and (5) assigns confidence classes to each transcript based on the evidence(s) that were used to construct those.

With a wide variety of available data for annotation, researchers often struggle to manage and optimize their usage. Several gene annotation software also offer users complicated configurations without providing substantial guidance. FINDER makes the job of gene annotation easy for bench scientists by automating the entire process from RNA-Seq data processing to gene prediction. Since FINDER does not assume the ploidy or the nucleotide composition of a genome, it could be applied to derive gene structures for a wide range of species, including non-model organisms. FINDER constructs gene models primarily from RNA-Seq data and is therefore capable of constructing tissue-and/or condition-specific isoforms which would have been impossible to obtain from ESTs only. FINDER supersedes the performance of existing software applications by utilizing read coverage information to fine-tune gene model boundaries. Instead of removing low-quality transcripts, FINDER flags them as low confidence – giving users the choice of using them as they seem fit. As a proof of concept, we provided evidence that using read coverage signal indeed enhances gene structures in a diverse set of organisms. Thus, we are confident that FINDER will pave the way for improved gene structure annotation in the future.

## Availability and requirements

Project name: FINDER

Project home page: https://github.com/sagnikbanerjee15/Finder

Operating system(s): Linux, MacOS

Programming language: Python, C, C++, Perl, Shell

License: MIT

Other software requirements: All software requirements are listed in https://github.com/sagnikbanerjee15/Finder/blob/master/environment.yml

### List of abbreviations

ESTs: Expressed Sequence Tags
NGS: Next Generation Sequencing
NCBI: National Center for Biotechnology Information
SRA: Sequence Read Archive
UTR: Untranslated Regions
CSV: Comma Separated Values
AED: Annotation Edit Distance
CPD: Changepoint Detection
TSS: Transcription Start Site
CDS: Coding Sequence
CPU: Central Processing Unit
cDNA: complementary DNA

## Declarations

### Ethics approval and consent to participate

Not applicable

### Consent for publication

Not applicable

### Availability of data and materials

FINDER can be accessed from https://github.com/sagnikbanerjee15/Finder RNA-Seq samples used for annotation is included in Additional file 8: Table S7 Barley PacBio sequences have been deposited in NCBI (Project id: GSE165730)

### Authors’ contributions

SB: Conceptualization, Data Curation, Formal Analysis, Investigation, Methodology, Software, Validation, Visualization, Writing – Original Draft Preparation, Writing – Review & Editing

PB: Formal Analysis, Writing – Review & Editing

MGW: Conceptualization, Resources, Supervision, Writing – Review & Editing

TZS: Conceptualization, Resources, Supervision, Writing – Review & Editing

RPW: Conceptualization, Investigation, Resources, Supervision, Writing – Review & Editing

CMA: Conceptualization, Funding Acquisition, Investigation, Project Administration, Resources, Supervision, Writing – Review & Editing

## Acknowledgements

This research used resources provided by the SCINet project of the USDA Agricultural Research Service, ARS project number 0500-00093-001-00-D. The authors are also thankful to Dr. Karin Dorman (Professor, Department of Statistics, Iowa State University), for providing insightful feedbacks to compare annotations and for implementing changepoint detection. The authors thank Gregory Fuerst for taking care of submitting data to NCBI and Dr. Eve Wurtele’s lab for the support in using Phylostratr.

## Additional Files

Additional file 1: Supplementary figures (S1-S9).

Additional file 2: Input to finder

Additional file 3: Annotation Edit Distance of reference transcripts as reported by each gene annotation pipeline

Additional file 4: Performance of gene annotation pipelines on coding regions of transcripts

Additional file 5: Comparison of FINDER’s performance with other gene annotation pipelines on a variety of different species

Additional file 6: Comparison of different transcriptome assembly softwares on a variety of species

Additional file 7: Improvement in reference gene annotation after adding untranslated regions verified with long-read from PacBio assemblies

Additional file 8: Description of RNA-Seq data used to execute FINDER, BRAKER2, MAKER2 and PASA

Additional file 9: Supplementary text document outlining methods and some results in more details

