## Additional file 1 for "FINDER: An automated software package to annotate eukaryotic genes from RNA-Seq data and associated protein sequences"

**Figure S1**

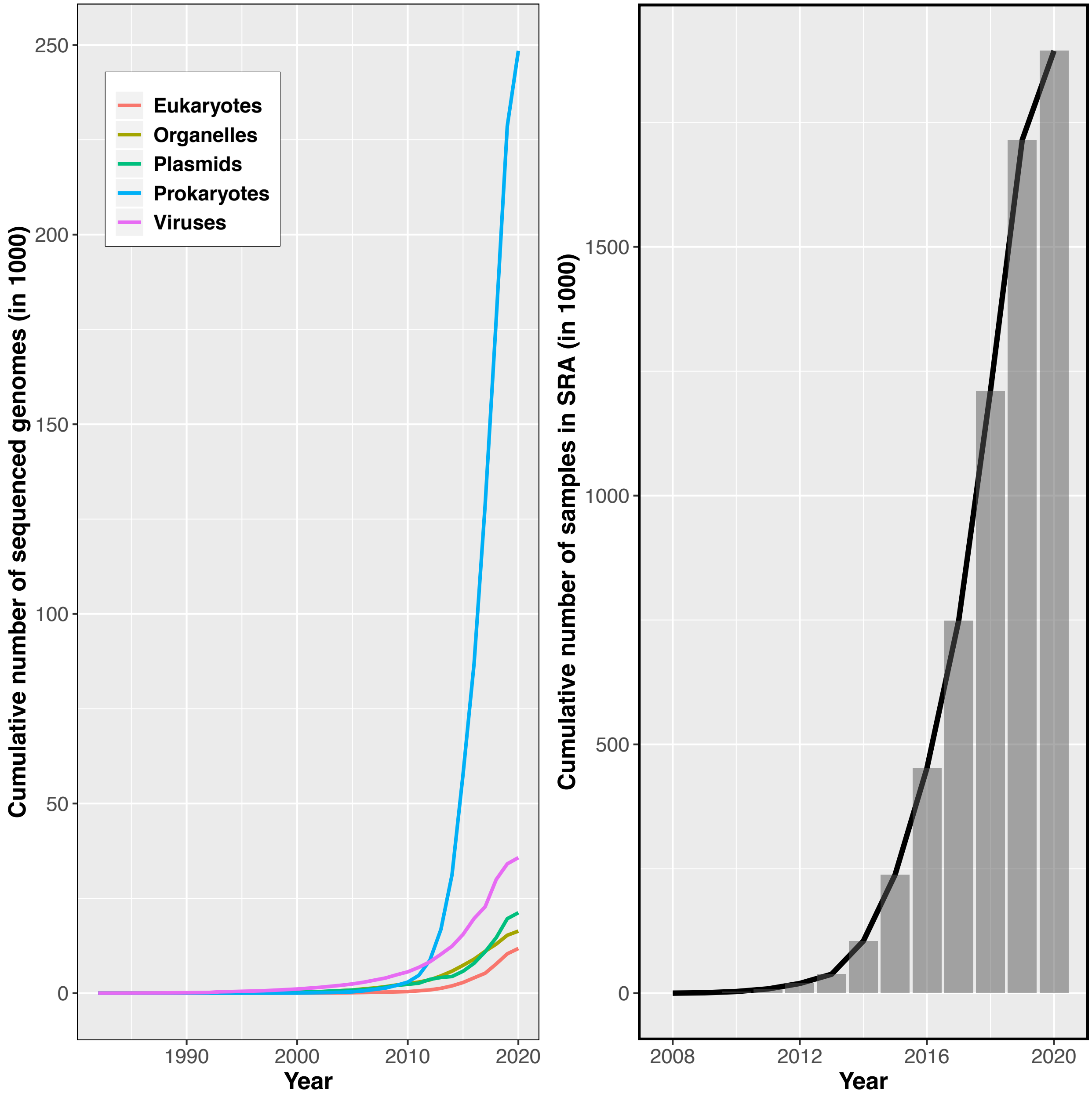


**Fig. S1** Increase in the number of genomes and the number of sequenced samples over the course of the last decade

#### Figure S2

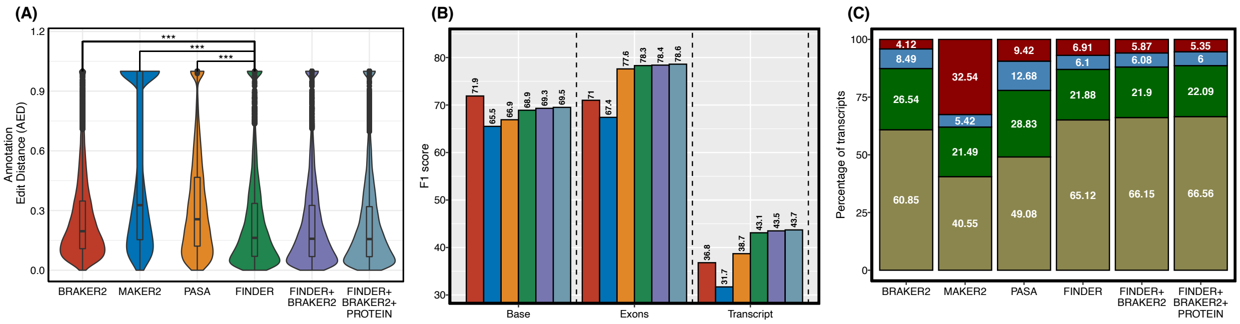


**Fig. S2** Performance comparison of predicted annotations with reference annotations of ***Drosophila melanogaster*** (A) Distribution of AED scores (B) Bar plot of F1 scores (C) Stacked bar plot showing percentage of transcripts in each of the four groups of AEDs

#### Figure S3

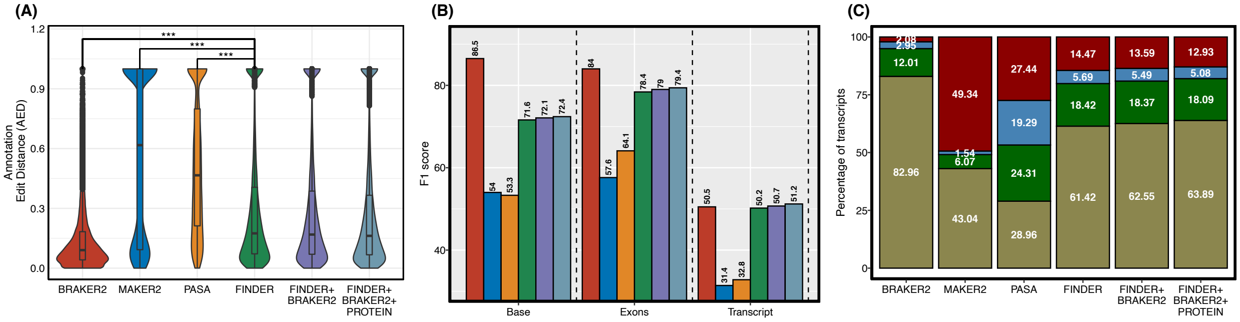


**Fig. S3** Performance comparison of predicted annotations with reference annotations of ***Caenorhabditis elegans*** (A) Distribution of AED scores (B) Bar plot of F1 scores (C) Stacked bar plot showing percentage of transcripts in each of the four groups of AEDs

### Figure S4

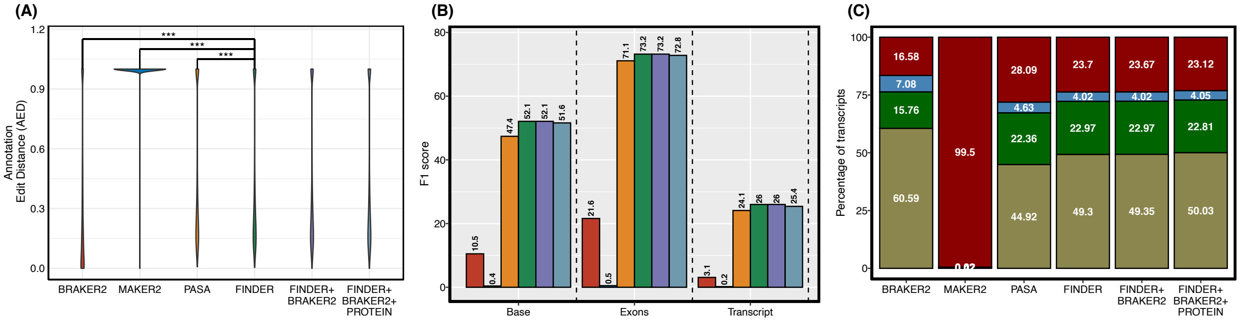


**Fig. S4** Performance comparison of predicted annotations with reference annotations of ***Hordeum vulgare*** (A) Distribution of AED scores (B) Bar plot of F1 scores (C) Stacked bar plot showing percentage of transcripts in each of the four groups of AEDs

### Figure S5

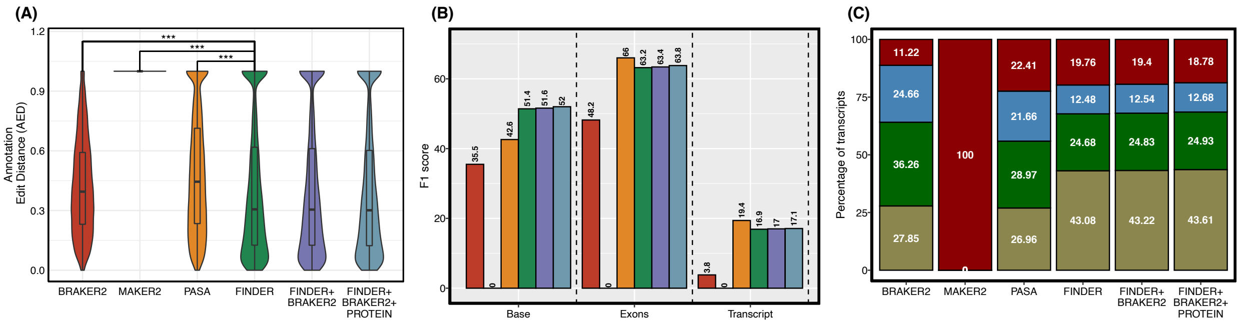


**Fig. S5** Performance comparison of predicted annotations with reference annotations of ***Homo sapiens*** (A) Distribution of AED scores (B) Bar plot of F1 scores (C) Stacked bar plot showing percentage of transcripts in each of the four groups of AEDs

#### Figure S6

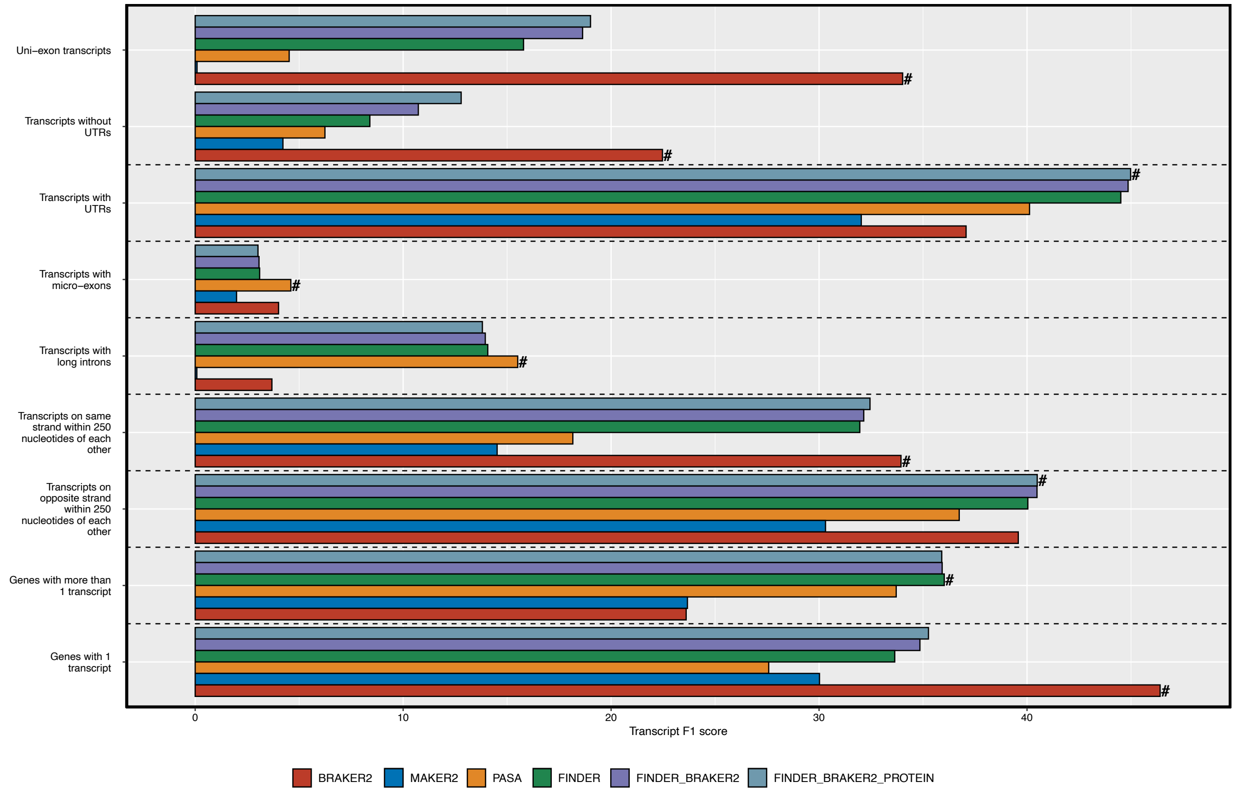


**Fig. S6** Comparison of performance of FINDER with other gene annotation pipelines on different groups of genes in *Drosophila melanogaster*

#### Figure S7

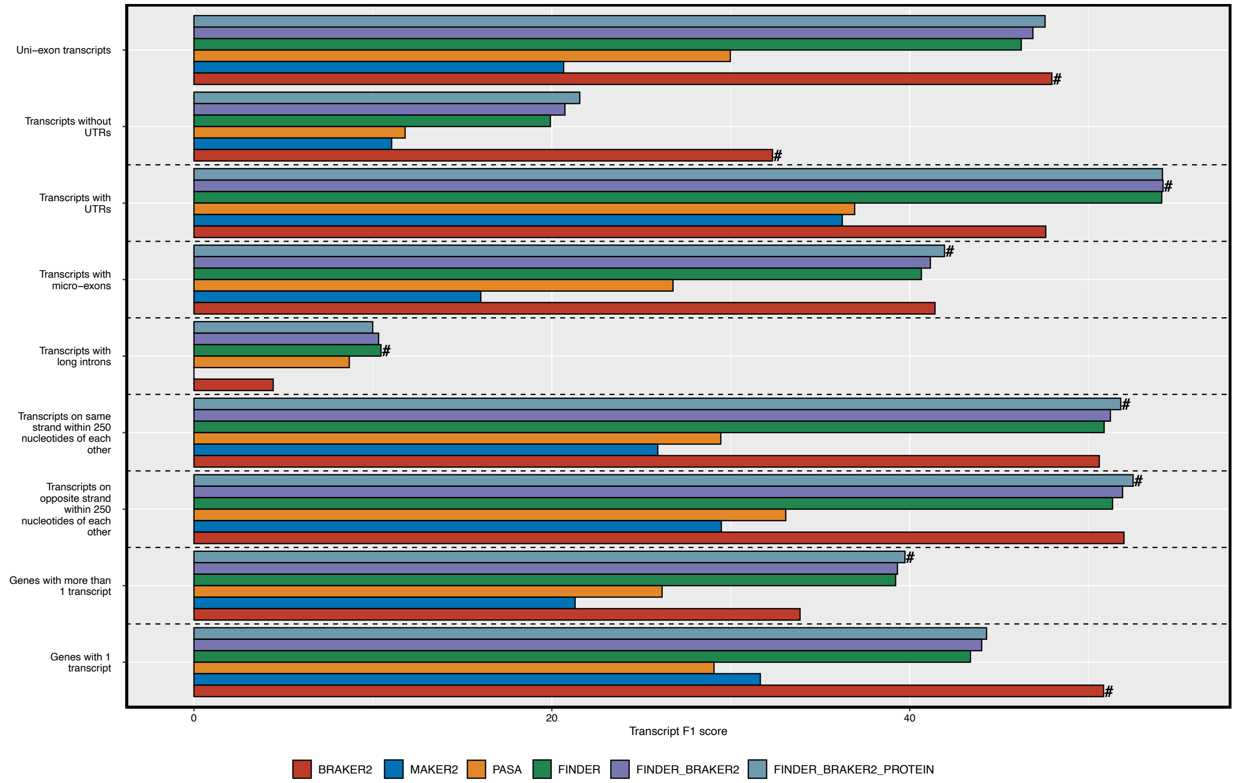


**Fig. S7** Comparison of performance of FINDER with other gene annotation pipelines on different groups of genes in *Caenorhabditis elegans*

#### Figure S8

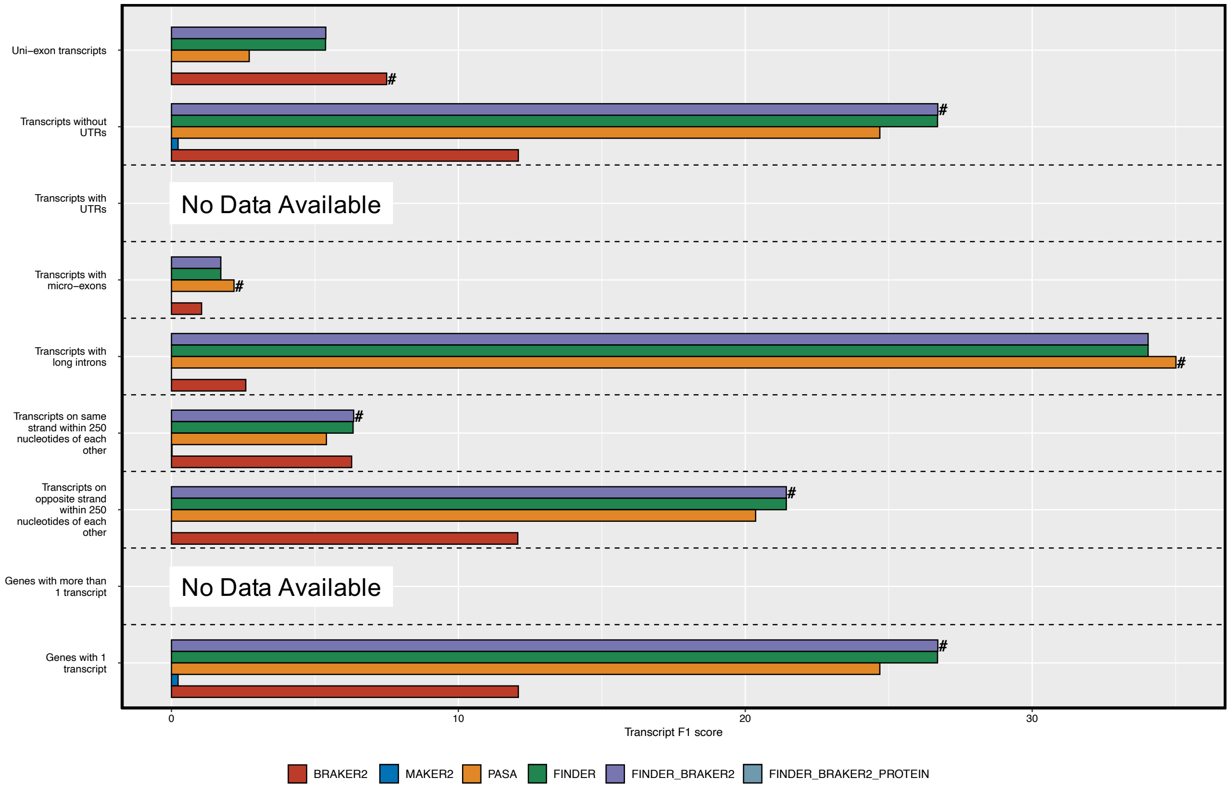


**Fig. S8** Comparison of performance of FINDER with other gene annotation pipelines on different groups of genes in *Hordeum vulgare*

#### Figure S9

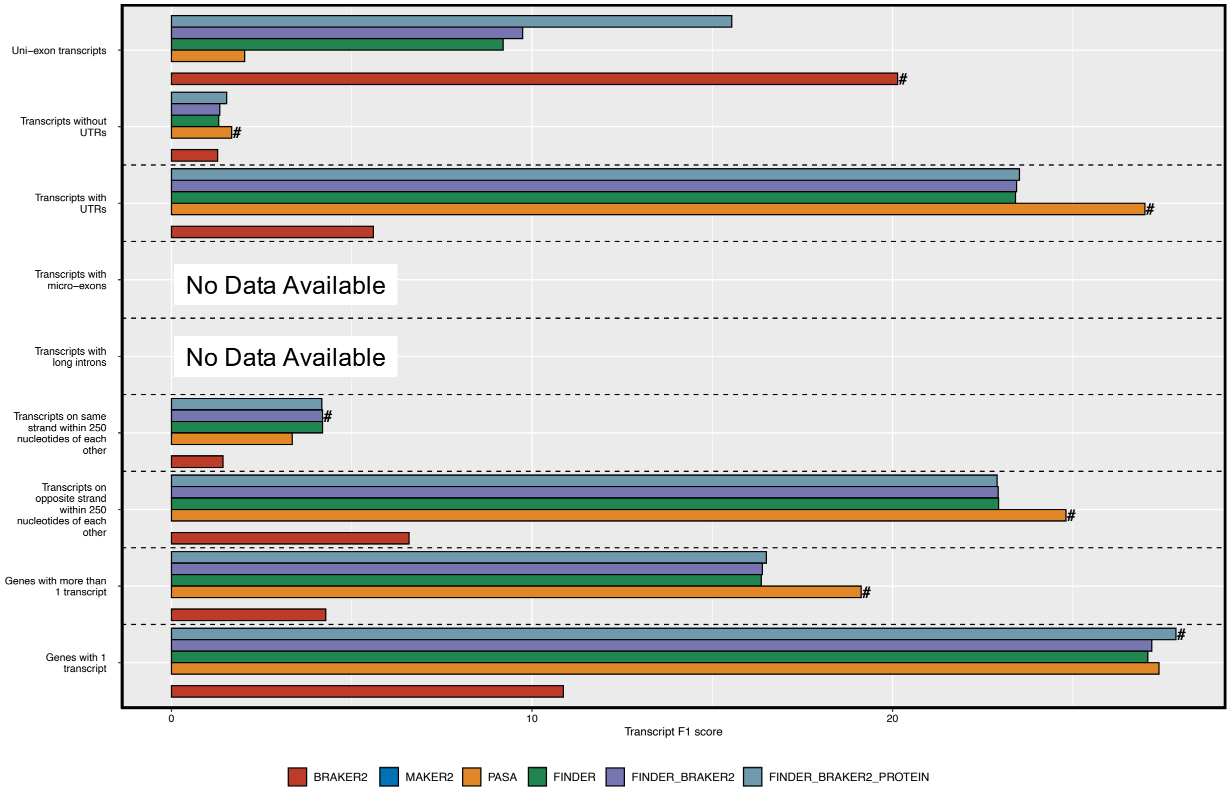


**Fig. S9** Comparison of performance of FINDER with other gene annotation pipelines on different groups of genes in *Homo sapiens*
