## Additional file 9 for "FINDER: An automated software package to annotate eukaryotic genes from RNA-Seq data and associated protein sequences"

Chapter 1

### Supplementary methods

FINDER is a gene annotator that uses RNA-Seq expression data to construct accurate gene structures with precise gene boundaries. In this chapter we have compiled the list of genomes and the reference gene annotations we used for the chosen species.

#### 1.1 Preparing data for testing

##### 1.1.1 Genome and reference gene annotations

We executed FINDER on 8 organisms. The genomes were downloaded from Ensemble. A mixture of small, medium and large genomes was used for testing.

##### 1.1.2 Expression data

FINDER was executed with RNA-Seq expression data downloaded from NCBI-SRA. Both single- and paired- ended read samples were used for the purposes of gene annotation. Read lengths varied from 75 bp to 600 bp. Samples sequenced only on illumine platform were used. Samples prepared by different techniques like oligo-DT, random priming, poly-T amplification, etc. were selected. For each organism, a variety of tissue type and conditions were chosen to approach an exhaustive transcriptome assembly. All details are provided in the Supplementary file @SRAData.

#### 1.2 Repeat masking

FINDER requires genomes to be soft masked since it is a requirement for BRAKER2 (Bruna et al. 2020; Hoff et al. 2016) to generate predicted gene models. FINDER's alignment and assembly modules do not require genomes to be soft masked. For testing, we used soft-masked genome sequences downloaded from Ensemble (**Error! Reference source not found.**)

#### 1.3 Alignments

FINDER is configured to run both STAR (Dobin et al. 2016, 2013) and OLego (Wu et al. 2013). STAR can align a very large number of reads in a short time and is highly configurable. FINDER runs multiple rounds of STAR with different settings to ensure optimal alignment. In the first pass, STAR only maps reads with a minimum overhang of 12 nucleotides, not allowing any soft clipping or mismatches. A filtering step selects splice junctions adequately supported by short reads and detected across multiple samples. Splice junctions are removed if the junction is present in less than four samples and is supported by fewer than three unique reads (or six multi-mapped reads) for canonical junctions, and seven unique reads (or ten multi-mapped reads) for non-canonical junctions in each of the samples. A cap of three samples have been chosen since it is likely for most experiments to have at least three replicates for a tissue type or condition. This highly stringent sifting procedure allows for selecting the most confident splice junctions supported by a sufficiently large number of perfectly mapped reads. FINDER compares read support across different samples of the same tissue type and/or condition. This is done with an attempt to preserve as much transcript diversity as possible.

The splice junction database generated in the first pass is used as an argument to the second pass. In the second pass, the alignment is relaxed to allow a mismatch of two nucleotides and a soft clipping of up to 5% of the read length. The minimum overhang was set to 8 nucleotides for splice junctions supported by the database and kept at 12 for novel splice junctions. Well supported splice junctions expressed across multiple samples of the same tissue type and/or condition are selected. A third pass is conducted to allow alignments to annotated splice junctions with the number of maximum mismatches increased to three. This step allows the read support for junctions to elevate without creating any new spurious splice junctions. The minimum and the maximum intron sizes for the first three rounds are set at 20 and 10,000 respectively. Several genes in plant cereal genomes have introns longer than 10kb (Li et al. 2009). To capture read alignments to large introns, STAR is executed for the final time with the minimum and the maximum intron sizes set at 10k and 10 million respectively. Outputs from each pass are combined to form a single set of alignments.

STAR focusses more on mapping reads faster but ignores those which arise from micro-exons. If STAR is allowed to map reads with large soft-clips (~50% of the length), then reads that arise from micro-exons will be mapped to it with soft-clips and cannot be used for assembling. Hence, we use OLego to map reads that are left unmapped by STAR. Unlike STAR, OLego focusses perfecting the alignments of short reads and takes more time to finish. Hence, FINDER only allows RNA-Seq samples having unmapped reads fewer than a million to be processed by OLego.

FINDER uses a total of 4 rounds of STAR to align reads. The first two rounds of mapping are intentionally kept stringent to allow best alignments and select those splice junctions that are supported by a large number of reads. Before moving on to the 3rd round of iteration, FINDER checks the percentage of reads aligned by STAR in the first two rounds. If fewer than 20% of reads are aligned, then FINDER reruns STAR will default settings to allow for most reads to map. Finally, read alignments from all the steps are merged into one by Samtools (Hoff et al. 2016; Bruna et al. 2020). The entire process has been summarized in Figure 1.1.

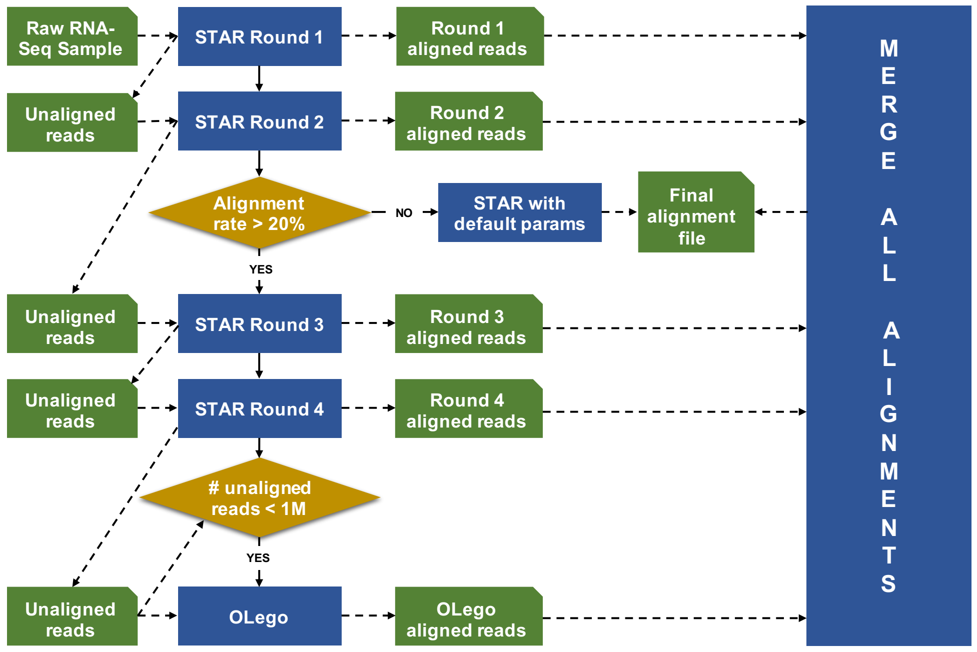

**Figure 1.1 Alignment Flowchart:** Step-by-Step protocol outlining the alignment process undertaken by FINDER.

#### 1.4 Assembling

PsiCLASS is a meta-aligner that generates a single transcriptome assembly from multiple RNA-Seq sources. It builds a global sub-exon graph by incorporating alignment information from all the samples, uses a mixture of gamma distributions to check if an exon is produced due to noise and deploys dynamic programming optimization to build an exhaustive and accurate set of transcripts.

On inspection of the output from PsiCLASS we noticed that the exon boundaries of several transcripts were truncated Figure 1.2. We consulted with the developers and modified PsiCLASS to extend the boundaries of the external exons to better accommodate the extent of RNA-Seq coverage (https://github.com/splicebox/PsiCLASS/issues/1).

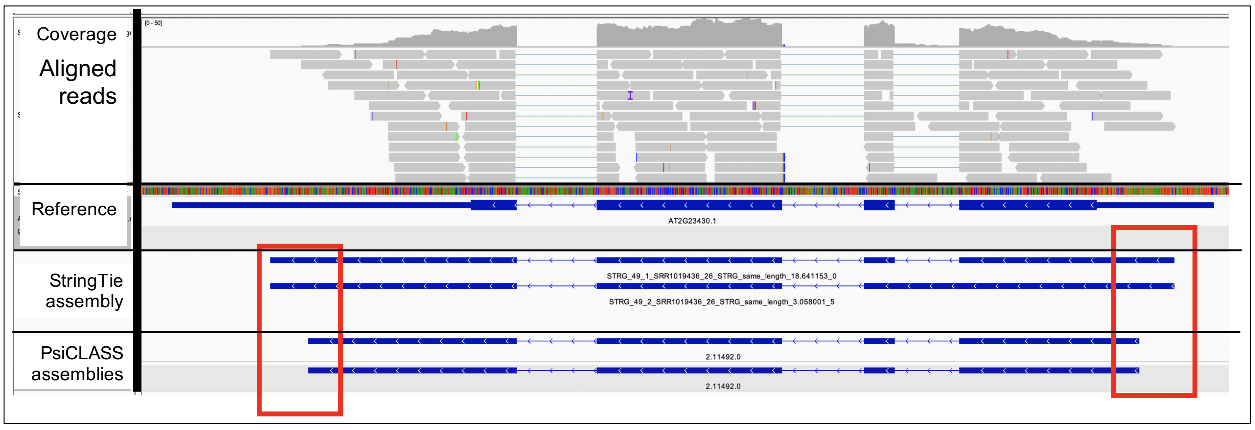

**Figure 1.2 Gene models generated by PsiCLASS have truncated external exons**

#### 1.5 Associating transcript models with conditions

PsiCLASS generates a meta-assembly from all the provided RNA-Seq alignments and also generates individual assemblies from each aligned file. PsiCLASS is designed to ignore gene models that appear in a small subset of samples since they are assumed to be spurious. FINDER scans each individual assembly and selects those gene models that were discarded by PsiCLASS. This allows FINDER to recognize transcripts that were expressed uniquely in specific tissues and/or conditions.

#### 1.6 Utilities included with FINDER

In addition to annotating genes in eukaryotes, `finder` offers 2 utilities that have a more generalized usage.

##### 1.6.1 verifyInputsToFINDER

Users can check if the inputs provided are compatible with `finder` or not using this utility. This program will scan the SRAdb database to ensure that the samples provided are indeed an RNA-Seq sample from the organism of choice. The program requires the path to the metadata file, the SRAmetadb file and the taxonomic id of the organism. It will report the samples that are not from an RNA-Seq sample or are from an incorrect taxon. We recommend users to perform a validation round using this utility to prevent the generation of incorrect annotations that can negatively impact downstream analysis.

##### 1.6.2 downloadAndDumpFastq

The downloadAndDumpFastq utility can used to download data from NCBI-SRA. It is optimized to use as many CPUs as requested by user thereby utilizing time optimally. It will also convert the SRA files to fastq files and remove the SRA files once the data download is complete. The program is configured to rerun the steps to ensure all the requested files have been downloaded.

Chapter 2

### Comparison with other gene annotation pipelines

We compared the gene annotations produced by FINDER with those generated by MAKER2 (Holt and Yandell 2011; Campbell et al. 2014), PASA (Haas et al. 2003) and BRAKER (Bruna et al. 2020; Hoff et al. 2016). This chapter outlines the series of commands used to generate gene annotations. We further provide more explanation about the analysis and results we reported in the main manuscript.

#### 2.1 Running BRAKER2

BRAKER2 offers several different options to predict gene annotations. We provided a soft masked genome and the aligned short reads as input to BRAKER2 (Figure 2 in https://github.com/Gaius-Augustus/BRAKER). We attempted to provide protein sequences as input to BRAKER2, but for most of the organisms the execution resulted in failure. Hence, we aligned protein sequences to the genome using exonerate (Slater and Birney 2005) and integrated it with the gene annotations produced from RNA-Seq short reads. BRAKER2 was not designed to utilize a large number of cores. So, FINDER was configured to use only 40 CPUs even when more resources were available. Also, BRAKER2 was unable to process genome fasta files when chromosome headers contained spaces. Genome fasta file headers were modified before providing them as inputs to BRAKER2. BRAKER2 is optimized to predict CDS but UTRs are added from RNA-Seq alignment files. We intended to compare RNA-seq evidence-based annotations only with prediction approaches. Hence, BRAKER2 predictions were generated without any UTRs.

BRAKER2 was executed using the following command:

```
braker.pl --genome <Path to organism genome> --cores <40 or less> --softmasking --overwrite --gff3 --bam
<list of comma separated bamfiles>
```

Alignment files were generated by FINDER according to the scheme outlined in the main manuscript.

#### 2.2 Assembling the transcriptome

Both PASA and MAKER2 requires assembled transcripts. We converted the final gtf transcriptome assembly reported by PsiCLASS to fasta format using gffread (Korf 2004).

```
gffread <gtf filename from PsiCLASS> -g <genome file name> -w <fasta filename>
```

We intentionally provided transcriptome assemblies generated by PsiCLASS (Korf 2004; Song et al. 2019) to ensure fairness of comparison, since FINDER uses the same for annotation. PsiCLASS was executed with the `--bamGroup` option turned on so that it can preserve features that are unique to a particular tissue/condition.

#### 2.3 Running PASA

To execute PASA two configuration files were generated using the following command:

```
echo "## templated variables to be replaced exist as <__var_name__>

# database settings
DATABASE=/project/maizegdb/sagnik/FINDER1_manuscript/data/finder1_runs/Arabidopsis_thaliana/pasa_psiclass/
arath.sqlite

#######################################################
# Parameters to specify to specific scripts in pipeline
# assign a value as done above.

#script validate_alignments_in_db.dbi
validate_alignments_in_db.dbi:--MIN_PERCENT_ALIGNED=<__MIN_PERCENT_ALIGNED__>
validate_alignments_in_db.dbi:--MIN_AVG_PER_ID=<__MIN_AVG_PER_ID__>

#script subcluster_builder.dbi
subcluster_builder.dbi:-m=50" > alignAssembly.config.txt

echo "## templated variables to be replaced exist as <__var_name__>

# database settings
DATABASE=/project/maizegdb/sagnik/FINDER1_manuscript/data/finder1_runs/Arabidopsis_thaliana/pasa_psiclass/
arath.sqlite

#######################################################
# Parameters to specify to specific scripts in pipeline
# assign a value as done above.

#script cDNA_annotation_comparer.dbi
cDNA_annotation_comparer.dbi:--MIN_PERCENT_OVERLAP=<__MIN_PERCENT_OVERLAP__>
cDNA_annotation_comparer.dbi:--MIN_PERCENT_PROT_CODING=<__MIN_PERCENT_PROT_CODING__>
cDNA_annotation_comparer.dbi:--MIN_PERID_PROT_COMPARE=<__MIN_PERID_PROT_COMPARE__>
cDNA_annotation_comparer.dbi:--MIN_PERCENT_LENGTH_FL_COMPARE=<__MIN_PERCENT_LENGTH_FL_COMPARE__>
cDNA_annotation_comparer.dbi:--MIN_PERCENT_LENGTH_NONFL_COMPARE=<__MIN_PERCENT_LENGTH_NONFL_COMPARE__>
cDNA_annotation_comparer.dbi:--MIN_FL_ORF_SIZE=<__MIN_FL_ORF_SIZE__>
cDNA_annotation_comparer.dbi:--MIN_PERCENT_ALIGN_LENGTH=<__MIN_PERCENT_ALIGN_LENGTH__>
cDNA_annotation_comparer.dbi:--MIN_PERCENT_OVERLAP_GENE_REPLACE=<__MIN_PERCENT_OVERLAP_GENE_REPLACE__>
cDNA_annotation_comparer.dbi:--
STOMP_HIGH_PERCENTAGE_OVERLAPPING_GENE=<__STOMP_HIGH_PERCENTAGE_OVERLAPPING_GENE__>
cDNA_annotation_comparer.dbi:--TRUST_FL_STATUS=<__TRUST_FL_STATUS__>
cDNA_annotation_comparer.dbi:--MAX_UTR_EXONS=<__MAX_UTR_EXONS__>
cDNA_annotation_comparer.dbi:--GENETIC_CODE=<__GENETIC_CODE__>" > annotCompare.config.txt
```

These configuration files were provided as input to the PASA pipeline.

```
Launch_PASA_pipeline.pl -c alignAssembly.config.txt \
-C \
--ALT_SPLICE \
--ALIGNERS gmap,blat \
--CPU 15 \
-R \
-g <Path to genome file> \
-t <Path to transcriptome fasta file generated by PsiCLASS> \
```

#### 2.4 Running MAKER2

MAKER2 is a gene annotator pipeline that uses both gene predictors and evidence from RNA-Seq experiments. We use SNAP (Stanke et al. 2008), AUGUSTUS (Tang et al. 2015) and GeneMark (Gremme et al. 2013) to generate predicted gene models. A total of three rounds of MAKER2 was executed. In each round, the genes predicted in the previous round was provided as input. A detailed step-by-step process has been provided below:

##### 2.4.1 Round1 MAKER2

```
# Round1 MAKER2 Run
maker -CTL # Generates configuration files
```

The file maker_opts.ctl was updated with the following:

```
genome=<Path to genome file>
est=<Path to transcriptome fasta file generated by PsiCLASS>
```

```
est2genome=1
cpus=<Number of CPUs requested by user>
```

Round1 of MAKER2 was launched with the following command:

```
maker -base <organism_name>
```

Annotations from all chromosomes were merged into one and then converted into gff3 using the following command:

```
gff3_merge -d <organism_name>_master_datastoreindex.log

maker2zff <organism_name>.all.gff3

mv <organism_name>.all.gff3 round1.gff3
```

Genometools (Venturini et al. 2018) was used to convert the gff3 file to gtf

```
gt gff3_to_gtf -force -o round1.gtf <(cat <(cat round1.gff|head -n +1) <(cat round1.gff|awk '$2=="maker"'))
```

In the next run, we adopted the method mentioned in the Supplementary document of Hoff 2016.

```
#Round2 MAKER2 run
cat round1.gtf | perl -ne '
    @t = split(/\t/);
    $seen{$t[8]} += ($t[4] - $t[3] + 1);
    if(eof()){
    $sum = 0; $c = 0;
    foreach my $key ( keys %seen ){
    $c=$c+1; $sum += $seen{$key};}
    print $sum."/".$c."=".($sum/$c);
    print "\n";
    }' > temp

flanking_region_length=temp/2
gff2gbSmallDNA.pl round1.gtf <genome file name> $flanking_region_length first.gb
```

##### 2.4.2 Training with SNAP

The following commands were executed to train SNAP

```
fathom -categorize 1000 genome.ann genome.dna
fathom -export 1000 -plus uni.ann uni.dna
forge export.ann export.dna
hmm-assembler.pl ${species} . > ${species}.hmm
```

##### 2.4.3 Training with AUGUSTUS

```
new_species.pl --species=${species}
etraining --species=${species} first.gb 1> etrain-test.out 2> etrain-test.err
fgrep "gene" etrain-test.err | cut -f 2 -d` ` > bad.etraining-test.lst
filterGenesOut_mRNAname.pl bad.etraining-test.lst first.gb > second.gb
etraining --species=${species} second.gb
optimize augustus.pl --species=$fspeciesg --onlytrain=second.gb.train.train second.gb.train.test
```

##### 2.4.4 Round2 MAKER2

Configuration files for the second round of MAKER2 run was generated using the following command.

```
maker -CTL # Generates configuration files
```

The file maker_opts.ctl was updated with the following:

```
genome=<Path to genome file>
est=<Path to transcriptome fasta file generated by PsiCLASS>
est2genome=1
```

```
cpus=<Number of CPUs requested by user>
maker_gff=round1.gff
est_pass=1
snaphmm=${species}.hmm # The hmm file generated after SNAP training
augustus=${species}
```

Round2 of MAKER2 was launched with the following command:

```
maker -base <organism_name>
```

##### 2.4.5 Training with GeneMark

GeneMark, another gene predictor, was used to generated predicted gene models using the following command:

```
gmes_petap.pl --ES --cores $CPU --sequence <genome file>
```

##### 2.4.6 Round3 MAKER2

For the final round, SNAP and AUGUSTUS was retrained using the steps outlined in the previous section. Configuration files for the third round of MAKER2 run was generated using the following command.

```
maker -CTL # Generates configuration files
```

The file maker_opts.ctl was updated with the following:

```
genome=<Path to genome file>
est=<Path to transcriptome fasta file generated by PsiCLASS>
est2genome=1
cpus=<Number of CPUs requested by user>
maker_gff=round2.gff
est_pass=1
snaphmm=${species}.hmm # The hmm file generated after SNAP training
augustus=${species}
gmhmm=${species}/gmhmm.mod
keep_preds=1
```

##### 2.4.7 MAKER2's performance

MAKER2's performance was the poorest for almost all organisms even though it was executed with three gene predictors and assembled RNA-Seq transcripts for three rounds. MAKER2 was able to report only a tiny fraction of the ground truth transcripts. This impacted the sensitivity which reduced the overall F1 score for MAKER2. Also, MAKER2's installation procedure was cumbersome and executing the program involved quite a bit of trial-and-error.

#### 2.5 Comparing FINDER gene models with other gene annotators

We used the `compare` utility from `mikado` (Venturini et al. 2018) and compared the gene annotations to the reference annotations obtained from Ensemble under the assumption that the reference annotations were gold standard and did not contain any errors. Mikado not only generates a summary of the comparison, but also provides comparison metrics for each individual transcript.

##### 2.5.1 F1 Score comparison

Mikado reports the number of transcripts in reference that is identified in the prediction and also the number of transcripts in the predicted annotation that have a perfect match with at least one reference transcripts. For multi-exonic transcripts, a perfect match is achieved when all the introns of the predicted transcript perfectly matches with the reference annotation. 2 mono-exonic transcripts form a perfect match only when at least 80% of the nucleotide definition agree with

one another. Mikado compare also reports agreement of the reference with the prediction in terms of nucleotides, exons and introns. We have used transcript matches, since they best represent the quality of annotation.
F1 score is computed as the harmonic mean between precision and recall. Recall is the fraction of reference transcripts that have a perfect match with at least one predicted transcript. Precision is the fraction of predicted transcripts that have a perfect match with at least one reference transcript. For an annotation to be good, both precision and recall should be high. While F1 score can indicate the number of transcripts correctly recognized, it does not offer any information about how the number of nucleotides in each reference transcript that was correctly annotated by the prediction. Hence, annotation edit distance has been used to determine how well each reference has been recognized.

##### 2.5.2 Annotation Edit Distance (AED) Score comparison

Annotation edit distance (AED) (Bao et al. 2013; Lu et al. 2013; Huang et al. 2016) is calculated using the following formula:

$$AED = 1 - \frac{2}{\left(\frac{1}{N\_precision} + \frac{1}{N\_recall}\right)}$$

Where,
N_precision = nucleotide level precision
N_recall = nucleotide level recall

Unlike F1-score, AED is reported for each reference transcript. A value of 0 indicates that the predicted transcript was a perfect match with the reference, while a score of 1 denotes that the reference was not reported. While constructing transcript from short read RNA-seq evidence it becomes quite a challenge to annotate the end exons correctly. This occurs due to low expression levels of some transcripts that may hinder the accurate reconstruction of the end points. Nonetheless, RNA-Seq data has sufficient information that could be leveraged to construct accurate untranslated regions (UTR). We selected those reference transcripts that had a perfect match in at least one of the reported gene annotations (BRAKER2, MAKER2, PASA or FINDER). AED scores were reported for only this set of transcripts. If a reference transcript, from this set, is not identified in a particular annotation then it is assigned a score of 1, specifically for that annotation. To further ensure that the AED scores were statistically lower for FINDER, we performed a Wilcoxon’s signed rank test. It is a non-parametric test that is widely used to assess whether the population mean ranks of two distributions differ significantly. In this case, we used the paired test, since each AED score corresponds to a particular reference transcript. All comparisons were made with FINDER reported transcripts.

#### 2.6 Processing long read sequencing data

As a proof-of-concept, we used PacBio sequences to show that FINDER produces better gene annotation. We downloaded the consensus FASTA sequence from six PacBio *Arabidopsis thaliana* transcriptome samples hosted in Isodb (Pertea et al. 2015; Kovaka et al. 2019) and aligned them to the *Arabidopsis thaliana* genome, using GMAP (Wu and Watanabe 2005), to generate a gff3 file. All sequences deposited in Isodb are error corrected with short reads from Illumina. These PacBio sequences are full-length transcripts, containing both CDS and UTRs, that were captured from RNA-Seq expression studies.

We performed the same analysis with *Hordeum vulgare* (Chapman et al. 2020; Hunt et al. 2019), since the r2 version of the gene annotation does not have any UTR annotation. We used RNA-Seq data generated in the Wise lab as documented in (Hunt et al. 2019) RNA-Seq transcripts, collected from blumeria treated barley leaves, were sequenced using 16 SMRT cells with replicates and size fractionations. Data was processed using the standard SMRTLink software version 6. Full-length transcripts were error corrected using HECIL (Choudhury et al. 2018), CoLoRMap (Haghshenas et al. 2016) and Hercules (Firtina et al. 2018).

Chapter 3

### Assemblers and assembly mergers

Improvements in technologies have resulted in a huge increase of sequencing experiments. This has necessitated the formulation of assembly softwares which are not only accurate but also very fast. Transcriptome assembly aided by alignments to the genome have shown to generate higher quality assemblies than *de novo* approaches (Liu and Dickerson 2017). There are several different genome-guided assemblers that are currently popular. Among them notable are Scallop (Shao and Kingsford 2017), Strawberry (Liu and Dickerson 2017), Stringtie (Pertea et al. 2015) and Trinity (Trapnell et al. 2012). We decided to skip Trinity since the genome guided approach defaults to the *de novo* approach and also it takes a very long time to complete.

#### 3.1 PsiCLASS assembler

All transcriptome assembly softwares take a single RNA-Seq aligned bamfile as an input and generates a single gtf assembly. PsiCLASS (Song et al. 2019) is a novel assembler that allows user to provide multiple RNA-Seq alignments as input. It constructs a consensus assembly for all the samples and also individual assemblies for each of the samples. PsiCLASS was executed with the `--bamGroup` option turned on indicating it to preserve all tissue/condition specific features.

#### 3.2 Other assemblers

To attest the supremacy of PsiCLASS over other assembly methods we compare it with Stringtie, Strawberry and Scallop. The arguments with which each assembler was executed has been provided below:

```
stringtie <alignment_filename> -p $CPU -o <output GTF filename>
scallop -i <alignment_filename> -o <output GTF filename>
strawberry --allow-multimapped-hits -p $CPU -o <output directory> <alignment filename>
```

StringTie is more sensitive to transcripts but also reports many false positives which reduces the overall transcript F1 score. Strawberry consistently predicts a large number of spurious transcripts for all the organisms, leading to a low specificity score. Scallop manages to attain average specificity and sensitivity. Both StringTie and Strawberry can be executed on multiple cores, but Scallop is designed for a single core machine. PsiCLASS achieves the best transcript F1 score by keeping the number of spurious transcripts at an absolute minimum thereby accomplishing the highest specificity score. At the same time, PsiCLASS also reports a sufficiently high fraction of the ground truth transcripts. Hence, FINDER uses only PsiCLASS to generate assemblies from short-read data.

#### 3.3 Merging assemblies together

Each of the assemblers we tested were able to generate a single GTF annotation file for each RNA-Seq sample. Hence, we needed to use other softwares to combine the GTF annotations from multiple RNA-Seq samples. We used Stringtie-merge (Pertea et al. 2015), cuffmerge (Niknafs et al. 2017; Trapnell et al. 2010), TACO (Niknafs et al. 2017) and Mikado (Venturini et al. 2018) to generate the consensus assemblies. All the softwares except Mikado completed the executed in under 5 minutes. Mikado took more than 15 hours to complete its execution and even then, it returned an empty gff3 file. Hence, we decided to eliminate Mikado from our study. We executed all the three merging softwares on *Arabidopsis thaliana*. As illustrated in Table 3.1, StringTie-merge generates the best transcriptome assembly registering an F1 score of 35.11 almost 10 units higher than cuffmerge. Hence, we used StringTie-merge for all the other organisms as well.

| | | Base Specificity | Base Sensitivity | Base F1 score | Exon Specificity | Exon Sensitivity | Exon F1 score | Transcript Specificity | Transcript Sensitivity | Transcript F1 score |
|---|---|---|---|---|---|---|---|---|---|---|
| **Stringtie** | *ST-merge* | 58.35 | 80.22 | 67.56 | 74.82 | 79.29 | 76.99 | 35.01 | 35.22 | **35.11** |
| | *TACO* | 75.58 | 69.53 | 72.43 | 88.19 | 71.28 | 78.84 | 25.39 | 24.47 | 24.91 |
| | *Cuffmerge* | 55.8 | 81.18 | 66.14 | 65.74 | 80.36 | 72.32 | 21.75 | 30.42 | 25.36 |
| **Scallop** | *ST-merge* | 37.34 | 85.3 | 51.94 | 70.64 | 79.67 | 74.88 | 24.84 | 32.19 | **28.04** |
| | *TACO* | 44.14 | 76.73 | 55.97 | 80.05 | 62.93 | 70.47 | 9.78 | 13.14 | 11.21 |
| | *Cuffmerge* | 36.45 | 86.55 | 51.29 | 66.19 | 78.68 | 71.89 | 17.15 | 28.8 | 21.49 |
| **Strawberry** | *ST-merge* | 38.41 | 87.06 | 53.3 | 43.86 | 85.3 | 57.93 | 6.88 | 31.68 | 11.3 |
| | *TACO* | 46.81 | 82.26 | 59.66 | 91.22 | 60.44 | 72.71 | 4.22 | 7.96 | 5.51 |
| | *Cuffmerge* | 41.76 | 84.75 | 55.95 | 53.54 | 79.13 | 63.87 | 8.28 | 25.45 | 12.49 |
| **PsiCLASS** | | 62.63 | 70.83 | 66.48 | 89.82 | 69.54 | 78.39 | 56.82 | 31.8 | **40.78** |
| **FINDER** | | 74.46 | 71.45 | 72.92 | 91.79 | 69.93 | 79.38 | 60.04 | 37.21 | **45.95** |

**Table 3.1 Comparison of performance of different softwares on combining multiple GTF annotations into a consensus annotation**
